# Neuronal NOX4 knockdown alleviates pathological tau-related alterations in a humanized mouse model of tauopathy

**DOI:** 10.1101/2020.10.14.338954

**Authors:** Enrique Luengo, Paula Trigo-Alonso, Cristina Fernández-Mendívil, Ángel Nuñez, Marta del Campo, César Porrero, Nuria García-Magro, Pilar Negredo, Cristina Sánchez-Ramos, Juan A. Bernal, Alberto Rábano, Jeroen Hoozemans, Ana I Casas, Harald H.H.W Schmidt, Ana María Cuervo, Manuela G. López

## Abstract

Approximately 44 million people worldwide live with Alzheimer’s disease (AD) or a related form of dementia. Aggregates of the microtubule-associated protein tau are a common marker of these neurodegenerative diseases collectively termed as tauopathies. However, all therapeutic attempts based on tau have failed, suggesting that tau may only indicate a higher-level causal mechanism. For example, increasing levels of reactive oxygen species (ROS) may trigger protein aggregation or modulate protein degradation. Here we show that type 4 NADPH oxidase (NOX), the most abundant isoform of the only dedicated reactive oxygen producing enzyme family, is upregulated in dementia and AD patients and in a humanized mouse model of tauopathy. Both global knockout and neuronal knockdown of the *Nox4* gene in mice, diminished the accumulation of pathological tau and positively modified established tauopathy by a mechanism that implicates modulation of the autophagy-lysosomal pathway (ALP). Moreover, neuronal-targeted NOX4 knockdown was sufficient to reduce neurotoxicity and prevented cognitive decline, suggesting a direct and causal role for neuronal NOX4. Thus, NOX4 is a previously unrecognized causal, mechanism-based target in tauopathies and blood-brain barrier permeable specific NOX4 inhibitors could have therapeutic potential even in established disease.

**Graphical abstract:** 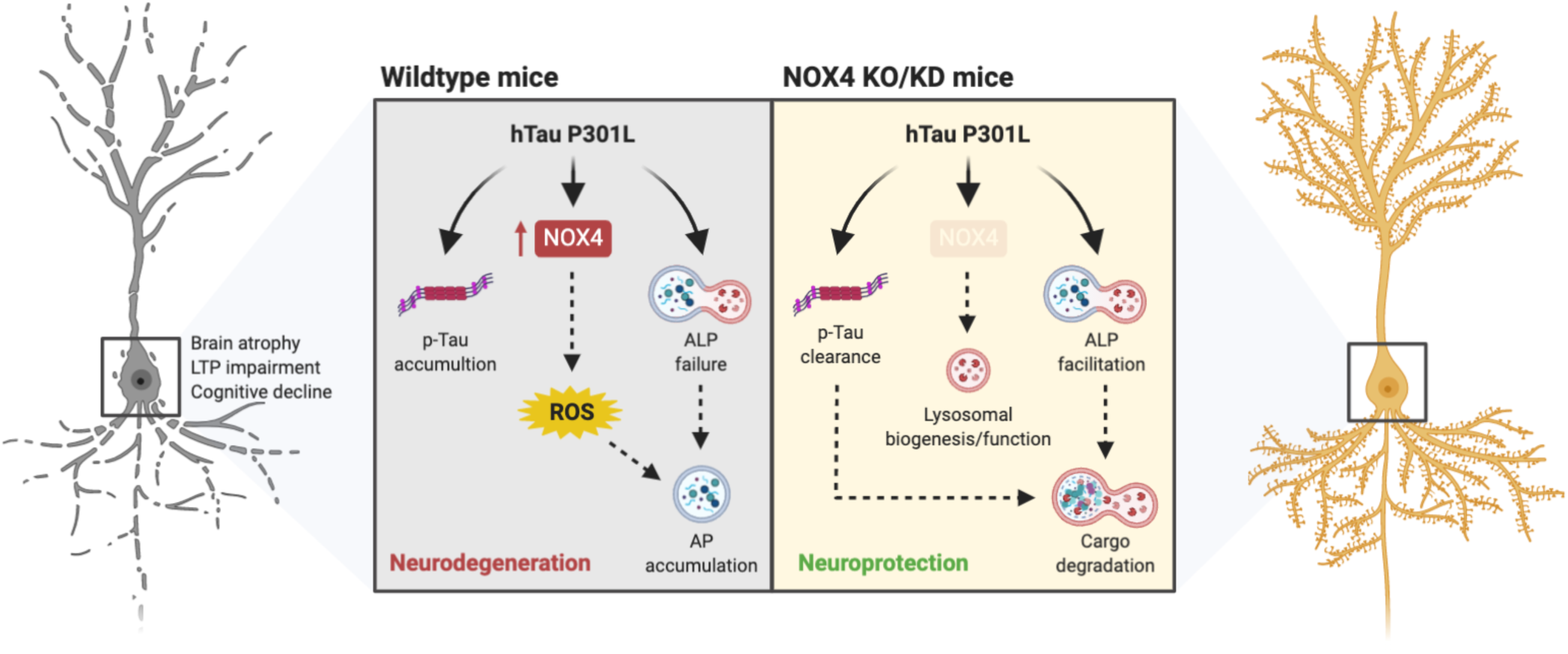

## Introduction

Major neurodegenerative disorders related to aging are considered proteinopathies as misfolded and aggregated proteins progressively accumulate, acquiring neurotoxic properties (1). In Alzheimer’s disease (AD), the most common form of dementia, histological hallmarks are represented by extracellular deposits of amyloid-β (Aβ) and intracellular aggregates of hyperphosphorylated microtubule-associated protein tau, known as neurofibrillary tangles (NFTs) (2, 3). Aggregates of tau protein characterize several neurodegenerative diseases (NDDs) that are collectively known as tauopathies, which include AD, frontotemporal dementia (FTD) and progressive supranuclear palsy (PSP), among others (2). Until now, there is no therapy available for AD despite the enormous human and economic efforts (4, 5). In recent years, therapeutic potential of targeting tau has received major attention, as tau pathology is more closely linked to cognitive decline than Aβ and shows good regional correspondence with neurodegeneration (2, 4, 5).

In relation to tau clearance, the ubiquitin-proteasome system and the autophagy-lysosomal pathway (ALP) are the major pathways involved in its removal (6). Among the different autophagy types, macroautophagy is a major intracytoplasmic protein degradation pathway characterized by the formation of double-membraned vesicles called autophagosomes that engulf cargo and target it to the lysosome for degradation and elimination of damaged organelles and large protein aggregates (7–9). Accumulating evidence supports ALP impairment in the pathogenesis of AD (1, 10, 11). ALP is upregulated in early stages of AD, but ALP-mediated clearance becomes impaired with disease progression, may promote autophagosome accumulation, inclusions of ubiquitinated proteins and neuritic dystrophy (1, 7, 12). Considering that tau is degraded via autophagy (6), dysfunction of ALP could lead to increased levels of its aggregated and toxic oligomeric forms (3, 13, 14). In addition, tau also inhibits autophagic degradation, disrupts autophagosome dynamics and induces lysosomal alterations, contributing to tau-induced toxicity (15–18) which strongly relates such to synaptic and cognitive deficits (19). All these evidences suggest that once tau aggregates are present in neurons, pathology can become self-perpetuating (3). Thus, given the involvement of defective autophagy in the pathogenesis and progression of tauopathies such as AD or FTD, therapies based on autophagy regulation can be implemented as ALP modulation accelerates degradation of tau protein aggregates (10, 20–22).

Reactive oxygen species (ROS) constitute one of the known autophagy regulators (23–26). NADPH oxidase (NOX), are considered the major enzymatic sources of ROS production (27, 28). Thus, NOX-derived ROS have been reported to regulate autophagy under different cellular conditions (26, 29). Among the members of this seven-member family (NOX1-5, DUOX1-2), NOX4, one of the main isoforms expressed in the central nervous system (CNS) (30), was found to regulate autophagy in energy-deprived cells (31). Moreover, NOX4 may be a key participant in the increased NOXs activity reported in AD progression as its expression is significantly increased in the brain of aged humanized APPxPS1 double transgenic mice (32). In line with this, we have recently demonstrated that neuronal NOX4 is a major contributor to cellular autotoxicity upon ischemia or hypoxia (33). In addition, neuronal NOX4 seems to play a pivotal role in progression of pathologies such as Parkinson’s disease (PD) or Traumatic Brain Injury (TBI) (34–36), and several studies have reported the involvement of NOXs family in cognitive dysfunction observed in AD patients and in *in vivo* AD models (32, 37, 38). Thus, while NOX4 has been involved in Aβ-related AD models (32, 39, 40) its implication with tau protein, the main driver of toxicity in AD, is currently unclear. Here, we aim to determine the potential implication of NOX4 in tau pathology and whether NOX4 could be validated as a novel therapeutic strategy for the treatment of tauopathies.

## Results

### NOX4 is overexpressed in tauopathy

To assess potential variations in the transcriptional profile of NOX isoforms in the context of tauopathy, we measured mRNA levels on postmortem brain tissue of Frontotemporal lobar degeneration (FTLD) and AD patients. In FTLD patients, only mRNA levels of NOX4 significantly increased by 2.8-fold (Figure 1A, left panel). In contrast, both *NOX2* (3.2-fold) and *NOX4* (2.2-fold) mRNA were significantly increased in AD patients compared to non-demented subjects (Figure 1A, right panel). These results demonstrate that *NOX4* mRNA levels are upregulated in both types of tauopathies, while *NOX2* mRNA levels are only augmented in AD. To better understand whether NOX4 was differentially overexpressed in distinct brain areas, we measured NOX4 protein levels in the hippocampus and prefrontal cortex, regions that are known to be remarkably affected during the progression of tauopathies. We observed that NOX4 was significantly increased in the hippocampus and prefrontal cortex of both FTLD (3.1 and 2.1-fold respectively) and AD patients (2.4 and 1.5-fold respectively) compared to non-demented subjects (Figure 1, B and C). Moreover, NOX4 protein levels correlated with mRNA levels, which were significantly increased in hippocampus and prefrontal cortex of both type of tauopathies (Supplemental Figure 1). These changes were further supported by immunofluorescence analysis of fixed postmortem brain sections that revealed increased levels of NOX4 in AT8 (pSer202/Thr205) positive hippocampal neurons from FTLD and AD patients (Figure 1D). To get more insight into the forms of tau that were more abundant in FTLD and AD patients in which NOX4 was overexpressed, we measured the oligomers (from 75 to 250 kDa) and monomers (from 50 to 65 kDa) (41, 42) of pathological hyperphosphorylated tau (Figure 1E). As observed, AT8 sarkosyl-insoluble (SI) oligomers and monomers were significantly augmented in FTLD (Figure 1F) and AD (Figure 1G) patients compared to non-demented subjects. However, no significant changes were observed in AT8 sarkosyl-soluble (SS) forms (Supplemental Figure 2). These results demonstrate an altered expression pattern of NOX4 in different brain areas of FTLD and AD patients in which insoluble forms of hyperphosphorylated tau are enriched, delineating a potential association between NOX4 and tau pathology.

**Figure 1:**
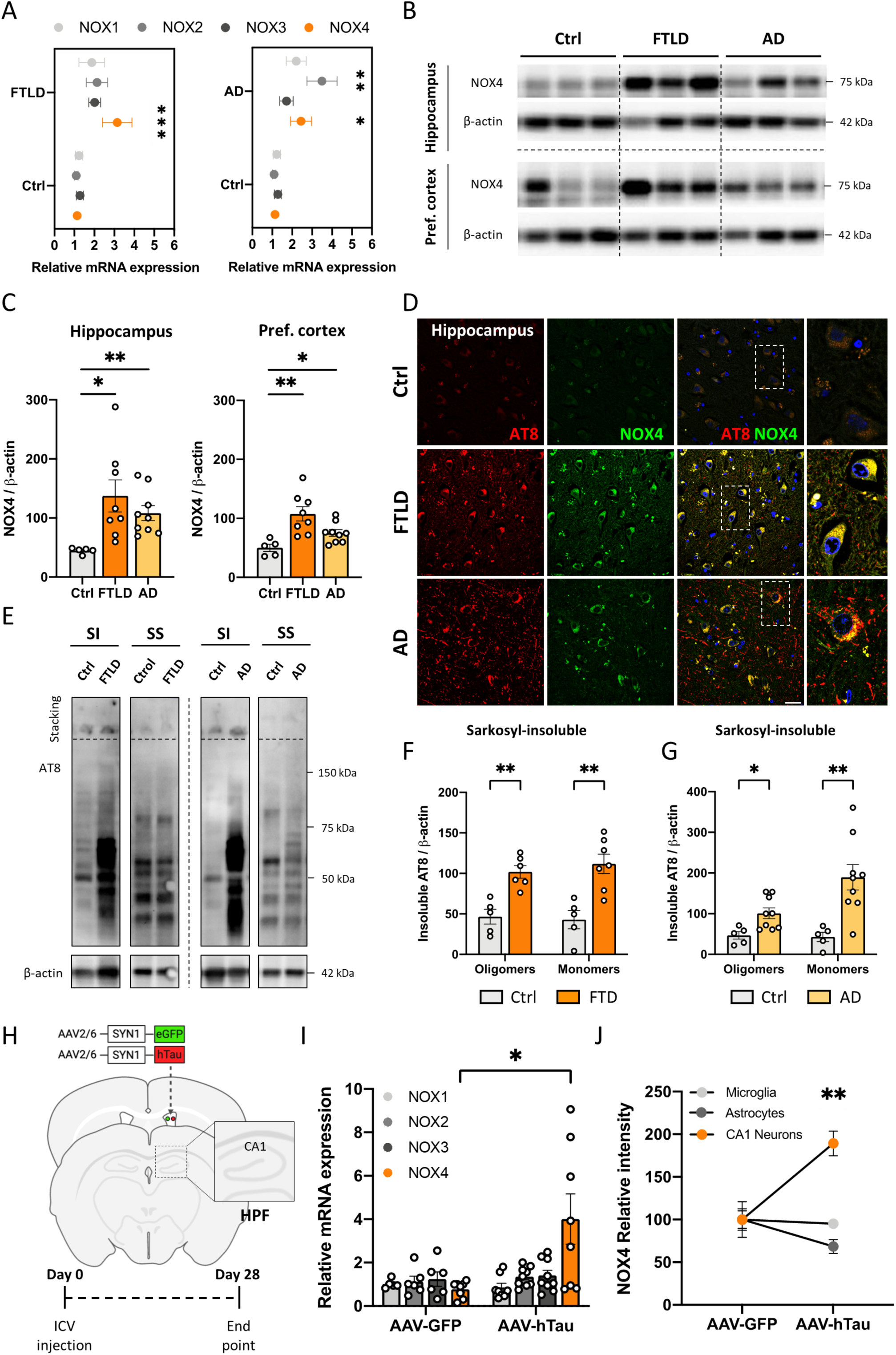
NOX4 is overexpressed in brains of FTLD and AD patients and in a humanized *in vivo* tauopathy model. **(A)** mRNA levels of NOX isoforms in postmortem human brain lysates from FTLD (left) (n=8), AD (right) (n=9) and non-demented subjects (Ctrl) (n=5). Representative images **(B)** and quantification **(C)** of NOX4 protein levels in hippocampus (up) and prefrontal cortex (down) from FTLD (n=8), AD (n=9) and Ctrl subjects (n=5). **(D)** Representative images of AT8 (p-tau S202/T205) and NOX4 intensity in fixed postmortem hippocampal sections of FTLD (n=2), AD (n=2) and Ctrl subjects (n=2). Insets show images at a higher magnification. Scale bar: 25 μm. 3 fields of view were acquired per subject. **(E)** Representative images of AT8 SI and SS tau oligomers and monomers from hippocampal lysates of FTLD, AD and Ctrl subjects. **(F)** Quantification of AT8 SI tau oligomers and monomers from FTLD (n=6-7) and Ctrl subjects (n=5). **(G)** Quantification of AT8 SI tau oligomers and monomers from AD (n=9) and Ctrl subjects (n=5). **(H)** Schematic representation of the protocol. (**I**) mRNA levels of NOX isoforms in hippocampal lysates from AAV-GFP (n=5-8) and AAV-hTau (n=8-9) injected mice. **(J)** NOX4 relative intensity in hippocampal CA1 neurons, microglia and astrocytes from AAV-GFP (n=3-6) and AAV-hTau (n=3-7) injected mice. Data are presented as mean ± SEM. Significance was determined by an unpaired Student’s t test or Mann-Whitney test for nonparametric data sets. *p<0.05; **p<0.01; ***p<0.001. SI (sarkosyl-insoluble); SS (sarkosyl-soluble).

To correlate the human results in an *in vivo* model that would allow us to study the implication of NOX4 in tau pathology, we used a well-characterized sporadic tauopathy model in mice (53). This model, consisted in the intracerebroventricular injection (ICV) of adeno-associated viral (AAV) vectors containing the human tau (hTau) mutation P301L (AAV-hTau), known to cause autosomal-dominant FTLD (43), or a control injection (AAV-GFP) in the right hemisphere’s ventricle. 28 days after AAV injection, behavioral and electrophysiological tests were performed and, thereafter, mice were sacrificed and the ipsilateral hippocampus was extracted (Figure 1H). While 28 days post-injection, a significant increase in hippocampal *NOX4* mRNA in AAV-hTau mice (5.2-fold) was observed, no significant differences were found in mRNA levels of other NOX isoforms (Figure 1I). Immunofluorescence analysis of fixed brain sections revealed 1.9-fold increase of NOX4 in the hippocampus of AAV-hTau injected mice compared to AAV-GFP controls (Figure 1J). Remarkably, the augmented levels of NOX4 were predominantly observed in pyramidal neurons of the hippocampal regions (CA1, CA2, CA3 and DG) of AAV-hTau mice (Supplemental Figure 3). No significant differences in NOX4 expression were found in either microglia or astrocytes (Figure 1J and Supplemental Figure 3) in the hippocampus. These results corroborate those obtained in patients and suggest that NOX4 overexpression, secondary to the presence of hTau P301L, is restricted to the neuronal compartment.

### The absence of NOX4 diminishes the accumulation of pathological tau

Once demonstrated that NOX4 was overexpressed in FTLD and AD patients and in the humanized tauopathy mouse model, we investigated whether the elimination of NOX4 could modify the accumulation of pathologic hyperphosphorylated tau *in vivo*.

Thus, we performed ICV injections of AAV-GFP/hTau particles in NOX4 knockout mice (NOX4^-/-^) and in wild type controls (NOX4^+/+^) (Figure 2A). We found a significant increase in AT8 inmunoreactivity (1.9-fold) in the AAV-hTau mice compared to AAV-GFP injected mice. Interestingly, NOX4^-/-^ mice did not show heightened AT8 levels 28 days after AAV-hTau injection (Figure 2, B and C). To further investigate which forms of tau were mostly affected by NOX4 deletion, we performed a separation protocol on NOX4^+/+^ and NOX4^-/-^ hippocampal homogenates to distinguish among the SI and SS fractions and measure pathological hyperphosphorylated AT8 and AT180 (pThr231) tau oligomers and monomers. We found a significant increase in both, AT8 SI and SS oligomers in NOX4^+/+^ mice injected with AAV-hTau compared to controls; this increase was partially abolished in NOX4^-/-^ mice. However, no significant changes were observed in either SI or SS AT8 monomers (Figure 2, D-F). We also found an increase in AT180 SI and SS monomers and oligomers in NOX4^+/+^ mice injected with AAV-hTau; however, this increase was reduced in NOX4^-/-^ mice (Figures 2, G-I). Overall, these results indicate that the absence of NOX4 reduces the accumulation of hyperphosphorylated tau, being the pathological tau oligomers the most affected form.

**Figure 2:**
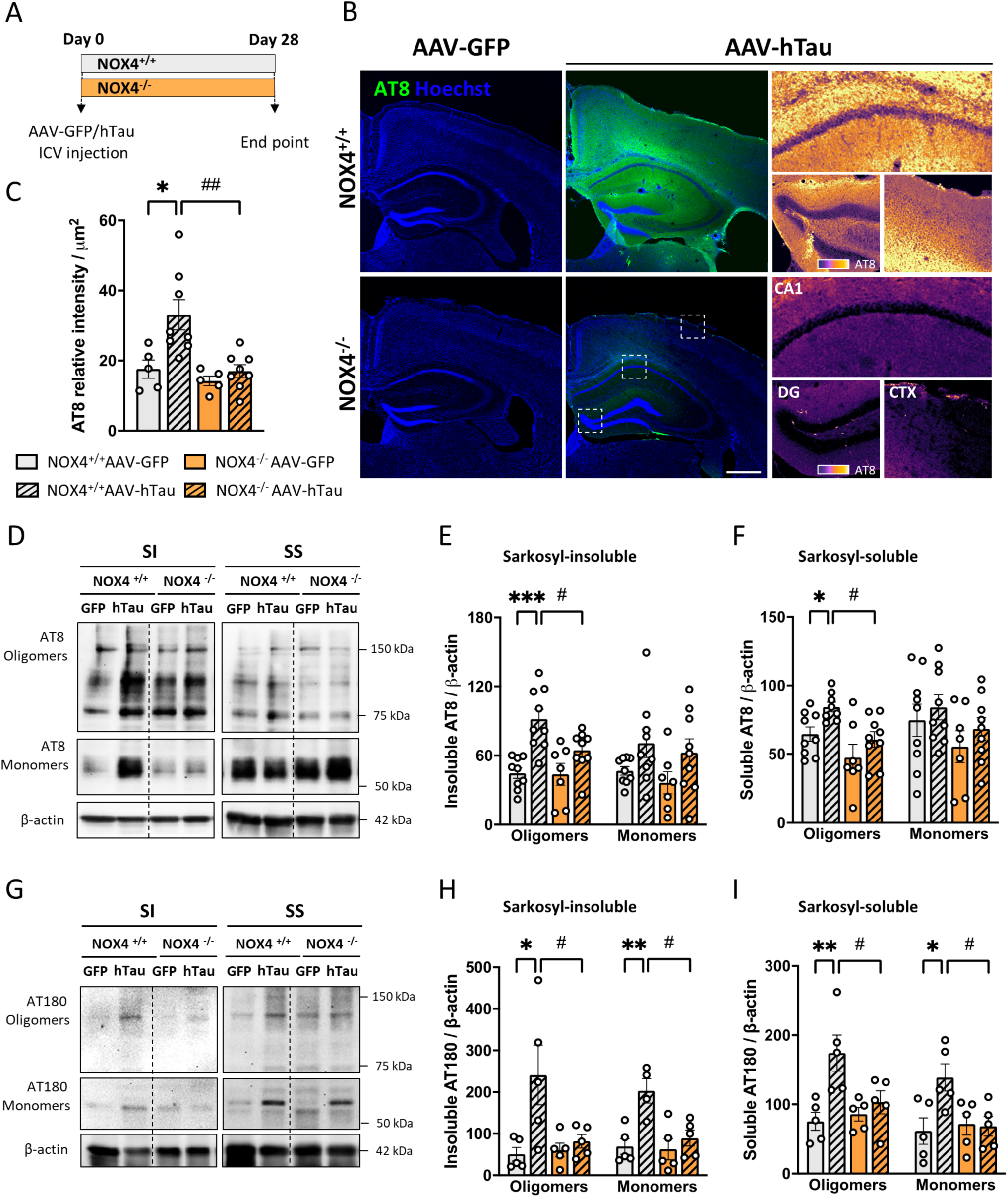
NOX4 genetic deletion reduces hyperphosphorylated tau accumulation 28 days after AAV-hTau injection. **(A)** Schematic representation of the protocol. Representative images **(B)** and quantification **(C)** of AT8 relative intensity in the hippocampus from NOX4^+/+^ (AAV-GFP, n=5 and AAV-hTau, n=8) and NOX4^-/-^ (AAV-GFP, n=5 and AAV-hTau, n=8) mice. Insets show images at a higher magnification. Scale bar: 500 μm (5X). Representative images **(D)** and quantification of AT8 SI **(E)** and SS **(F)** tau oligomers and monomers from hippocampal lysates of NOX4^+/+^ (AAV-GFP, n=9 and AAV-hTau, n=9) and NOX4^-/-^ (AAV-GFP, n=7 and AAV-hTau, n=8) mice. Representative images **(G)** and quantification of AT180 SI **(H)** and SS **(I)** tau oligomers and monomers from hippocampal lysates of NOX4^+/+^ (AAV-GFP, n=5 and AAV-hTau, n=5) and NOX4^-/-^ (AAV-GFP, n=5 and AAV-hTau, n=5) mice. Data are presented as mean ± SEM. Significance was determined by one-way ANOVA with Tukeýs post hoc test. *p<0.05; **p<0.01; ***p<0.001; #p<0.05; ##p<0.01. DG (Dentate Gyrus); CTX (Retrosplenial cortex).

### NOX4 deficiency modulates the ALP in tauopathy

A substantial amount of evidence supports that autophagy dysregulation occurs in AD patients and in AD animal models (44). Noteworthy, a large amount of immature autophagic vacuoles accumulate in dystrophic neurites in AD brains (45), suggesting that the accumulation of pathogenic proteins such as Aβ and tau may be caused by a defective ALP performance (46, 47). As NOX4 deficiency reduced the accumulation of pathological oligomeric hyperphosphorylated tau, we sought of interest to evaluate the performance of macroautophagy, a highly characterized proteolysis pathway involved in the clearance of tau (48, 49), in the *in vivo* tauopathy model. Immunofluorescence analysis revealed that the macroautophagy markers p62 (Figure 3, A and B) and microtubule-associated protein 1 light chain 3 (LC3) (Figure 3, C and D) were accumulating in the ipsilateral CA1 hippocampal region of NOX4^+/+^ AAV-hTau injected mice. As observed, NOX4 genetic deletion prevented this alteration in macroautophagy secondary to tau injection (Figure 3, A-D). These results were confirmed by western blot analysis of ipsilateral hippocampus lysates that showed increased expression of p62 and LC3 II in NOX4^+/+^ AAV-hTau but not in NOX4^-/-^AAV-hTau injected mice (Figure 3, E-G). However, in AAV-hTau injected mice, the absence of NOX4 did not changed p62 and LC3 mRNA levels, suggesting a post-transcriptional regulation of this clearance process (Supplemental Figure 4). These results suggest that NOX4 could be, at least in part, involved in tau-mediated macroautophagy dysregulation. To gain further insights on how NOX4 was regulating macroautophagy in tauopathy, we analyzed by immunofluorescence the nuclear translocation of Transcription Factor EB (TFEB), a key regulator of lysosomal biogenesis, and the immunoreactivity against its target gene, the lysosomal-associated membrane protein 1 (LAMP1). Our results revealed that nuclear localization of TFEB, together with LAMP1 expression, were increased in CA1 hippocampal neurons in NOX4^-/-^ AAV-hTau mice (Figure 3, H-J). These results were corroborated by western blot analysis in which we observed increased levels of LAMP1 and Cathepsin D (CTSD), another target gene of TFEB, in NOX4^-/-^ AAV-hTau compared to NOX4^+/+^ AAV-hTau injected mice (Figure 3G, K and L). Taken together, these results indicate that deletion of the *NOX4* gene in mice prevents the macroautophagy blockade secondary to hTau and increases the expression of lysosomal-related proteins, unveiling NOX4 as a potential modulator of lysosomal pathway.

**Figure 3:**
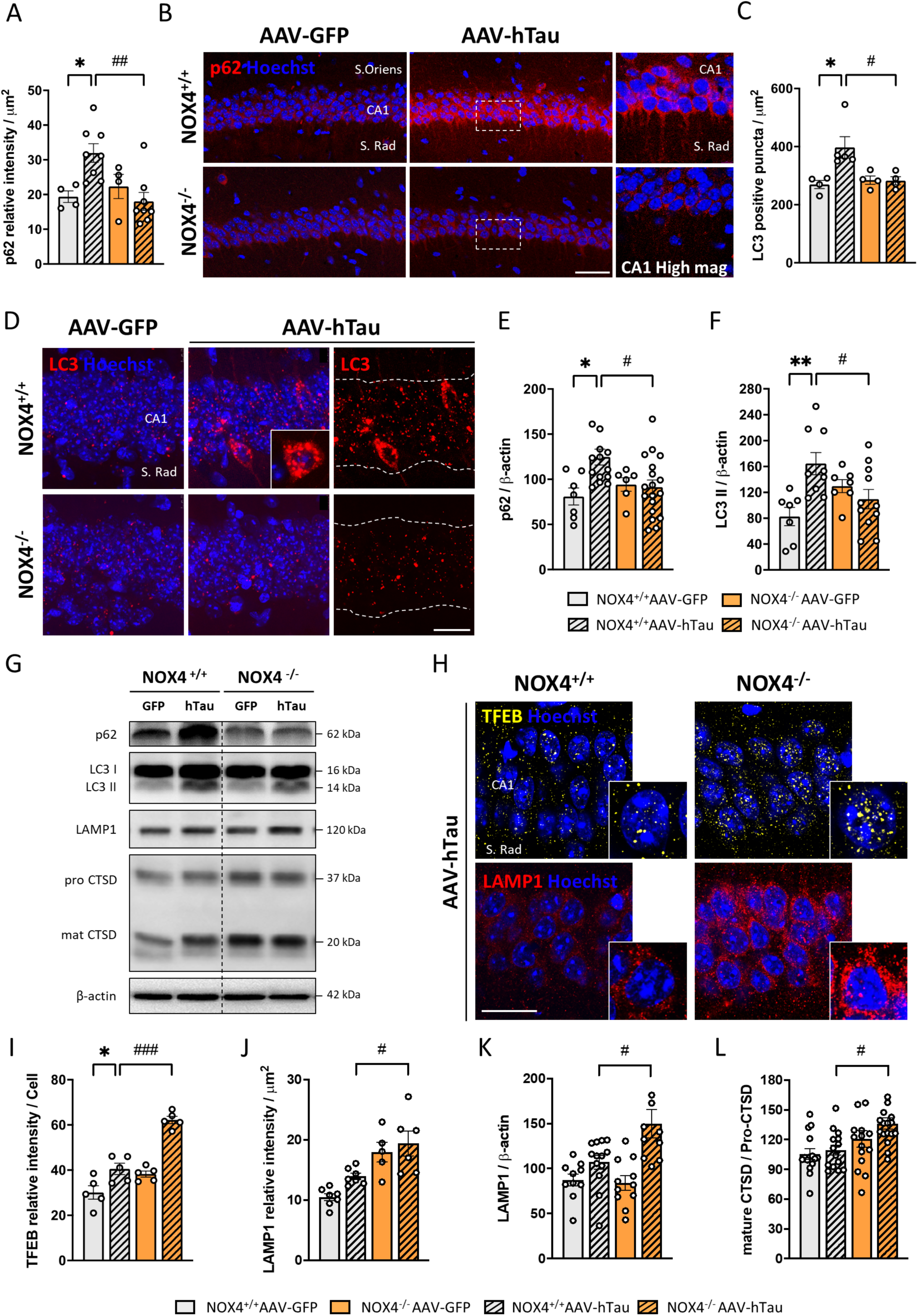
Macroautophagy blockade induced by AAV-hTau injection is prevented in NOX4 deficient mice. **(A)** Quantification and representative images **(B)** of p62 relative intensity in the hippocampal CA1 region from NOX4^+/+^ (AAV-GFP, n=4 and AAV-hTau, n=8) and NOX4^-/-^ (AAV-GFP, n=4 and AAV-hTau, n=8) mice. Insets show images at a higher magnification. Scale bar: 25 μm (40X). **(C)** Quantification and representative images **(D)** of LC3 positive puncta in the hippocampal CA1 region from NOX4^+/+^ (AAV-GFP, n=4 and AAV-hTau, n=5) and NOX4^-/-^ (AAV-GFP, n=4 and AAV-hTau, n=4) mice. Scale bar: 15 μm (63X). Quantification of p62 **(E)** and LC3 II **(F)** protein expression levels from hippocampal lysates of NOX4^+/+^ (AAV-GFP, n=7 and AAV-hTau, n=14) and NOX4^-/-^ (AAV-GFP, n=6 and AAV-hTau, n=18) mice. **(G)** Representative images of protein expression levels of macroautophagy and lysosomal markers from hippocampal lysates of NOX4^+/+^ and NOX4^-/-^ mice injected with AAV-GFP or AAV-hTau. **(H)** Representative images of nuclear TFEB and LAMP1 relative intensity in the hippocampal CA1 region from NOX4^+/+^ and NOX4^-/-^ mice injected with AAV-hTau. Insets show images at higher magnification. Scale bar: 15 μm (63X). Quantification of nuclear TFEB **(I)** and LAMP1 **(J)** relative intensity in the hippocampal CA1 region from NOX4^+/+^ (AAV-GFP, n=5-7 and AAV-hTau, n=5-7) and NOX4^-/-^ (AAV-GFP, n=5 and AAV-hTau, n=5-6) mice. Quantification of LAMP1 **(K)** and mature CTSD **(L)** protein expression levels from hippocampal lysates of NOX4^+/+^ (AAV-GFP, n=10-15 and AAV-hTau, n=13-14) and NOX4^-/-^ (AAV-GFP, n=11-15 and AAV-hTau, n=10-17) mice. Data are presented as mean ± SEM. Significance was determined by one-way ANOVA with Tukeýs post hoc test. *p<0.05; **p<0.01; #p<0.05; ##p<0.01; ###p<0.001. S.Oriens (Stratum Oriens); S.Rad (Stratum Radiatum); SI (sarkosyl-insoluble); SS (sarkosyl-soluble).

In addition to the protein aggregation/accumulation and protein clearance imbalance, oxidative stress and inflammation are known to play a pivotal role in the progression of tauopathies (50, 51). Furthermore, NOX-derived ROS are essential signals to regulate autophagy (52, 53). Thus, we assessed whether NOX4 genetic deletion could modulate ROS production and inflammation secondary to AAV-hTau injection (Supplemental Figure 5A and 6A). While in NOX4^+/+^ AAV-hTau injected mice ROS production was increased in the hippocampus, in NOX4^-/-^ AAV-hTau mice it was reduced to control levels (Supplemental Figure 5, B-D). Related to pro-inflammatory proteins, inducible nitric oxide synthase (iNOS), the microglial activation marker CD68 and the main components of the inflammasome complex (caspase-1 and IL-1β) were upregulated in hippocampal lysates of NOX4^+/+^ AAV-hTau compared to NOX4^+/+^ Control mice. These parameters were also remarkably reduced in NOX4^-/-^ AAV-hTau mice (Supplemental Figure 6, B-F). These results highlight a key role of NOX4 in modulating ROS production and neuroinflammation in the humanized *in vivo* tauopathy model.

**Figure 5:**
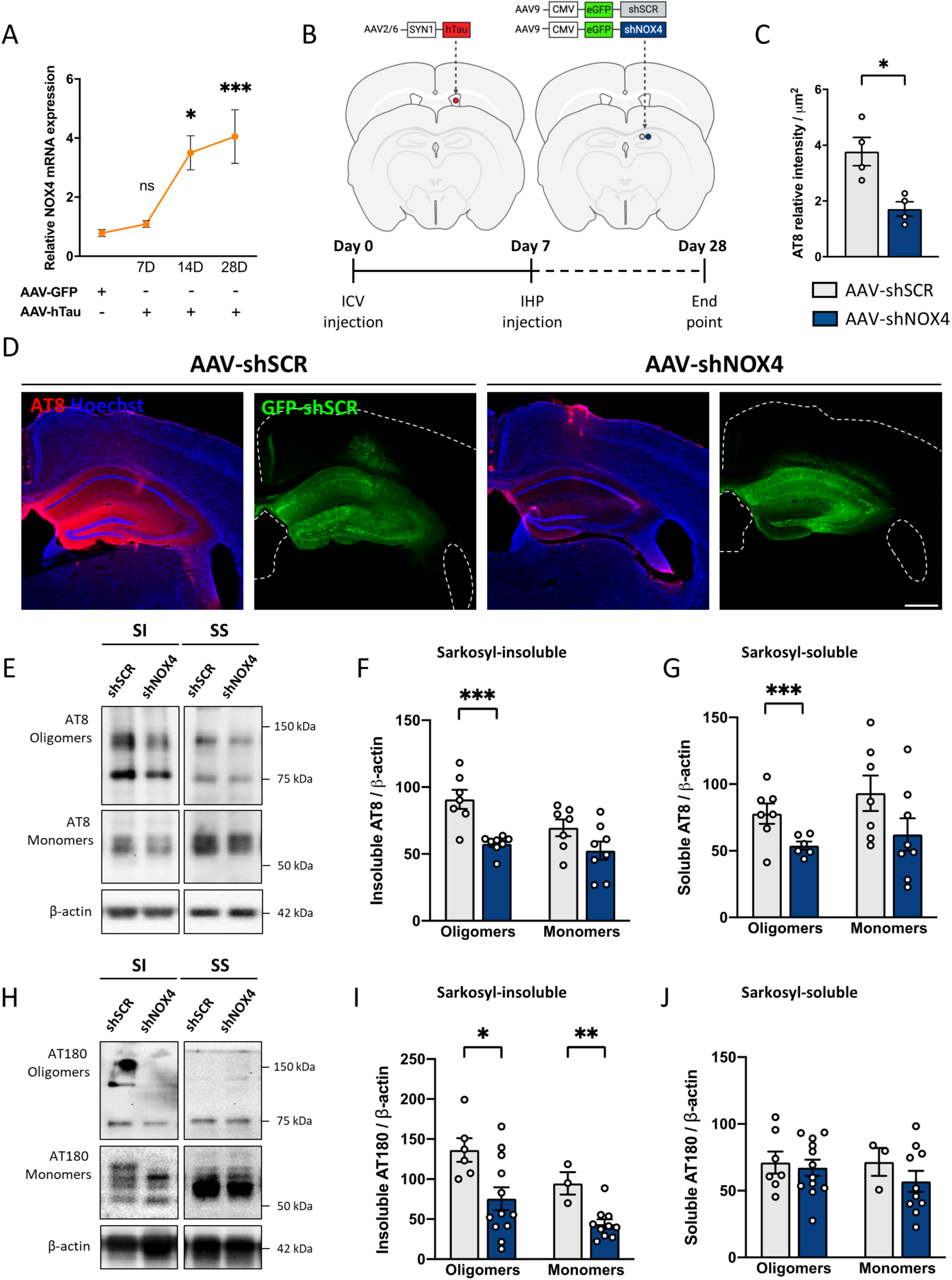
Neuronal-targeted NOX4 knockdown diminishes the accumulation of hyperphosphorylated tau once tauopathy is initiated. **(A)** *NOX4* mRNA from hippocampal lysates of NOX4^+/+^ (AAV-GFP, n=14 and AAV-hTau, n=8) and NOX4^-/-^ (AAV-GFP, n=4 and AAV-hTau, n=11) mice. **(B)** Schematic representation of the protocol. Quantification **(C)** and representative images **(D)** of AT8 relative intensity in the hippocampus of AAV-hTau (AAV-shSCR, n=4 and AAV-shNOX4, n=4) mice. Scale bar: 500 μm (5X). Representative images **(E)** and quantification of AT8 SI **(F)** and SS **(G)** tau oligomers and monomers from hippocampal lysates of AAV-hTau (AAV-shSCR, n=7 and AAV-shNOX4, n=6-8) mice. Representative images **(H)** and quantification of AT180 SI **(I)** and SS **(J)** tau oligomers and monomers from hippocampal lysates of AAV-hTau (AAV-shSCR, n=3-7 and AAV-shNOX4, n=10-12) mice. Data are presented as mean ± SEM. Significance was determined by one-way ANOVA with Tukeýs post hoc test or unpaired Student’s t test. *p<0.05; **p<0.01; ***p<0.001. SI (sarkosyl-insoluble); SS (sarkosyl-soluble).

**Figure 6:**
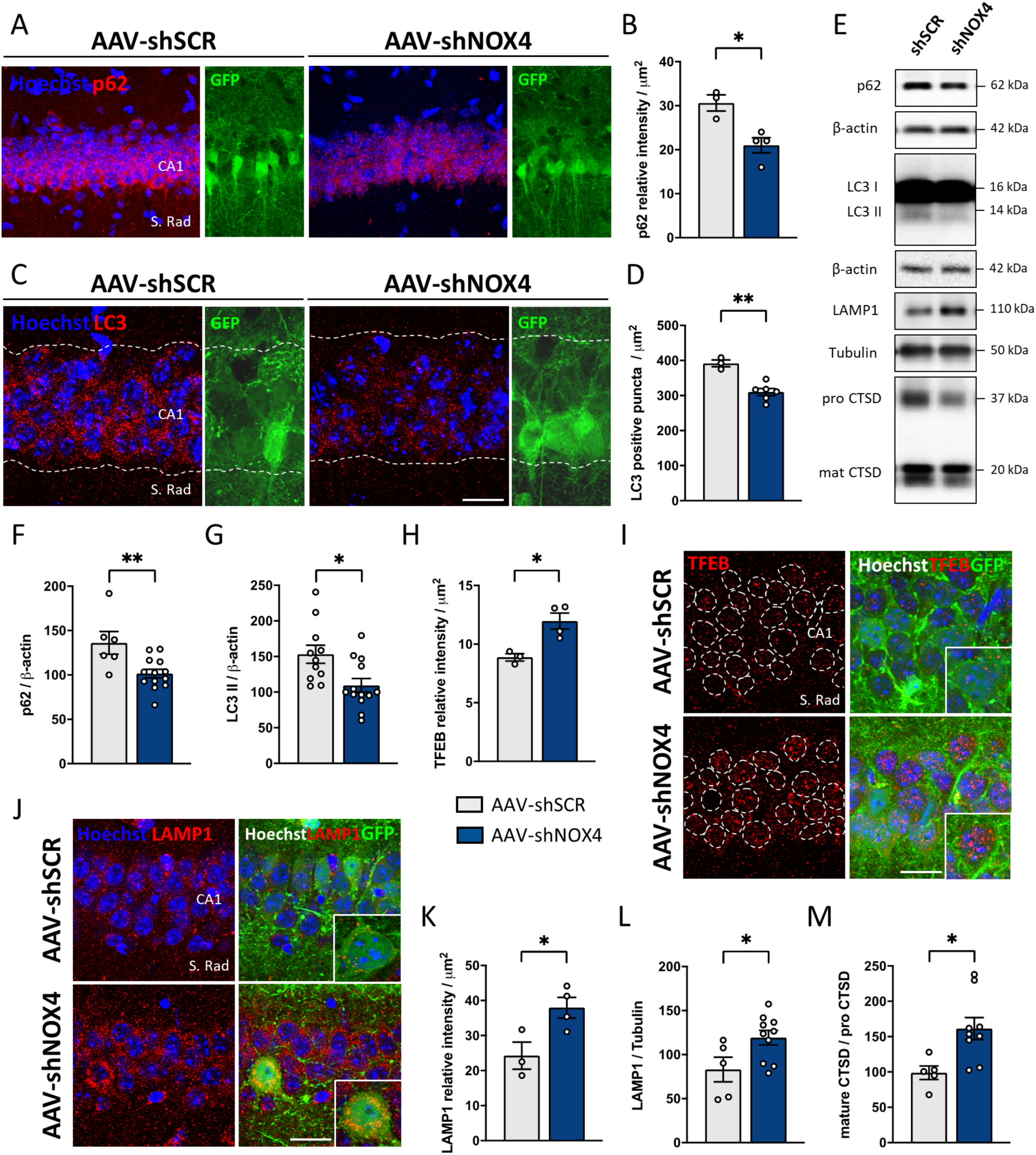
NOX4 neuronal knockdown prevents macroautophagy blockade once tauopathy is initiated. Representative images **(A)** and quantification **(B)** of p62 relative intensity in the hippocampal CA1 region from AAV-hTau (AAV-shSCR, n=3 and AAV-shNOX4, n=4) mice. Scale bars: 25 μm (40X). Representative images **(C)** and quantification **(D)** of LC3 positive puncta in the hippocampal CA1 region from AAV-hTau (AAV-shSCR, n=3 and AAV-shNOX4, n=6) mice. Scale bars: 15 μm (63X). **(E)** Representative images of protein expression levels of macroautophagy and lysosomal markers from hippocampal lysates of AAV-shSCR and AAV-shNOX4 injected mice. Quantification of p62 **(F)** and LC3 II **(G)** protein expression levels from hippocampal lysates of AAV-hTau (AAV-shSCR, n=6-11 and AAV-shNOX4, n=13) mice. Quantification **(H)** and representative images **(I)** of nuclear TFEB in the hippocampal CA1 region from AAV-hTau (AAV-shSCR, n=3 and AAV-shNOX4, n=4) mice. Scale bars: 7.5 μm (63X). Representative images **(J)** and quantification **(K)** of LAMP1 relative intensity in the hippocampal CA1 region from AAV-hTau (AAV-shSCR, n=3 and AAV-shNOX4, n=4) mice. Scale bars: 7.5 μm (63X). Quantification of LAMP1 **(L)** and CTSD **(M)** protein expression levels from hippocampal lysates of AAV-hTau (AAV-shSCR, n=5 and AAV-shNOX4, n=9-10) mice. Data are presented as mean ± SEM. Significance was determined by unpaired Student’s t test. *p<0.05; **p<0.01. S.Rad (Stratum Radiatum).

### NOX4 genetic deletion prevents functional and cognitive impairments in tauopathy

To determine whether NOX4 deficiency contributes to tau-related neuropathology, we measured the thickness of layers in regions typically affected during the course of tauopathies, such as cortex and hippocampus. The retrosplenial cortex (CTX), the pyramidal cell layer in CA1 region (CA1) and the top granule cell layer in the dentate gyrus (DG TOP), were noticeably and significantly thinner in NOX4^+/+^ AAV-hTau injected mice compared to NOX4^+/+^ AAV-GFP controls. Strikingly, the absence of NOX4 largely attenuated the cellular loss and the brain atrophy observed in the *in vivo* tauopathy model (Figure 4, A and B). Besides neuronal loss, pathogenic tau has shown to mediate synaptic dysfunction in different tauopathy/AD models (54–56). Thus, we investigated whether NOX4 genetic deletion could prevent tau-associated impairment of long-term potentiation (LTP) *in vivo*, a highly-characterized method to assess synaptic plasticity and a cellular model for learning and memory (57, 58). While NOX4^+/+^ and NOX4^-/-^ mice injected with AAV-GFP presented a sustained hippocampal LTP over 30 minutes, NOX4^+/+^ AAV-hTau exhibited LTP deficits. However, the absence of NOX4 alleviated LTP impairments secondary to hTau injection (Figure 4, C-E).

**Figure 4:**
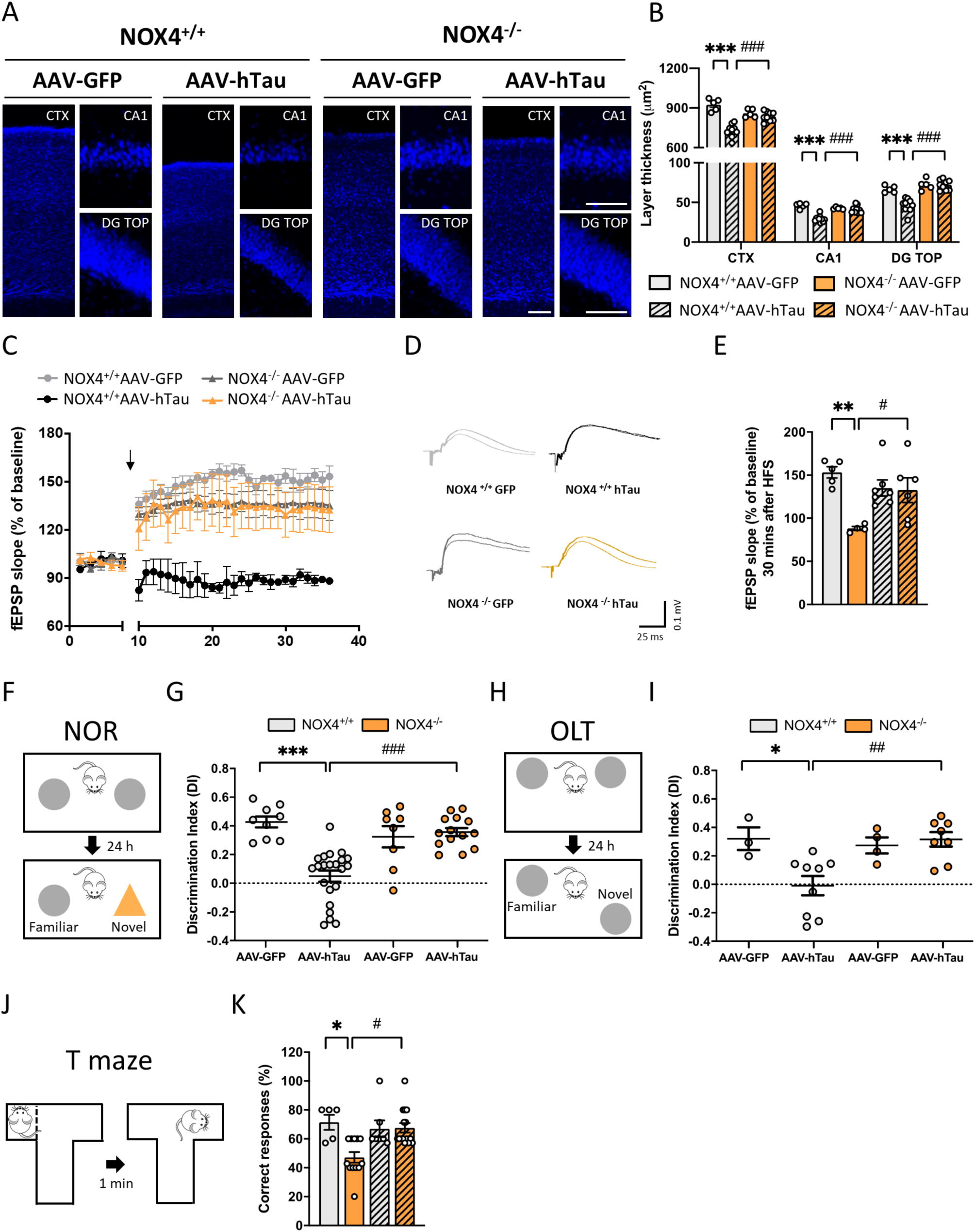
Brain atrophy, LTP impairment and cognitive decline triggered by AAV-hTau injection are reduced in NOX4^-/-^ mice. Representative images **(A)** and thickness quantification **(B)** of CTX, CA1 and DG TOP in NOX4^+/+^ (AAV-GFP, n=5 and AAV-hTau, n=10) and NOX4^-/-^ (AAV-GFP, n=5 and AAV-hTau, n=9) mice. Scale bars: 100 μm (10X); 75 μm (40X). **(C)** *In vivo* long-term potentiation (LTP) in NOX4^+/+^ and NOX4^-/-^ mice injected with AAV-GFP or AAV-hTau over 30 minutes. Arrow indicates high-frequency stimulation (HFS). Five minutes of control period and 30 minutes after HFS stimulation are shown. **(D)** Representative traces before and after HFS. **(E)** Quantification of the fEPSP slope 30 minutes after HFS in NOX4^+/+^ (AAV-GFP, n=5 and AAV-hTau, n=4) and NOX4^-/-^ (AAV-GFP, n=7 and AAV-hTau, n=6) mice. **(F)** Schematic representation of the NOR test. **(G)** Quantification of the DI in NOX4^+/+^ (AAV-GFP, n=9 and AAV-hTau, n=22) and NOX4^-/-^ (AAV-GFP, n=8 and AAV-hTau, n=14) mice. **(H)** Schematic representation of the OLT. **(I)** Quantification of the DI in NOX4^+/+^ (AAV-GFP, n=3 and AAV-hTau, n=9) and NOX4^-/-^ (AAV-GFP, n=4 and AAV-hTau, n=8) mice. **(J)** Schematic representation of the T maze. **(K)** Quantification of the percentage of correct responses in NOX4^+/+^ (AAV-GFP, n=5 and AAV-hTau, n=12) and NOX4^-/-^ (AAV-GFP, n=7 and AAV-hTau, n=15) mice. Data are presented as mean ± SEM. Significance was determined by one-way ANOVA with Tukeýs post hoc test or Kruskal-Wallis with Dunńs post hoc test for nonparametric data sets. *p<0.05; **p<0.01; ***p<0.001; #p<0.05; ##p<0.01; ###p<0.001. CTX (Retrosplenial cortex); DG TOP (Dentate gyrus top layer).

To assess whether NOX4 genetic deletion could restore tau-associated cognitive decline, we performed different behavioral tasks. First, the novel object recognition test (NOR) (Figure 4F) that assesses recognition memory (59, 60). Both, NOX4^+/+^ and NOX4^-/-^ mice injected with AAV-GFP exhibited preference for the novel object over the familiar one, displaying a normal cognition as indicated by the discrimination index (DI).

However, NOX4^+/+^ AAV-hTau injected mice spent equal amounts of time exploring both objects, resulting in clear cognitive impairment represented by a significant decrease in DI. On the contrary, NOX4^-/-^ AAV-hTau injected mice showed a performance similar to that of AAV-GFP control injected mice, indicating a sustained cognitive performance (Figure 4G). Next, hippocampus-dependent spatial memory and spatial learning were evaluated using the object location test (OLT) (Figure 4H) and the T maze (Figure 4J), respectively. In the OLT, similar results were achieved to those in the NOR test; NOX4^-/-^ AAV-hTau showed improved performance compared to NOX4^+/+^ AAV-hTau injected mice (Figure 4I). In agreement with these results, in the T maze test, NOX4^+/+^ AAV-hTau injected mice failed to recognize the unexplored arm during the test time, which is indicative of cognitive deficit (Figure 4K), whereas NOX4^-/-^ AAV-hTau mice exhibited a performance similar to that of control AAV-GFP injected mice, indicating an improvement on spatial memory (Figure 4K). Overall, these results demonstrate that the absence of NOX4 alleviates brain atrophy and prevents LTP impairment and cognitive decline induced by hTau P301L.

### Neuronal NOX4 knockdown efficiently reduces the accumulation of pathological *tau*

The positive results obtained in the NOX4^-/-^ mice, together with the observation that NOX4 overexpression was mainly restricted to the neuronal compartment upon hTau P301L injection, encouraged us to explore whether the protective effects of NOX4 deficiency against tauopathy were mainly attributed to neuronal NOX4. To address this, neuronal NOX4 knockdown in hippocampal neurons was performed 7 days after ICV AAV-hTau injection. At this time, *NOX4* mRNA was not yet increased (Figure 5A), and although mice did not present cognitive decline, tauopathy and its related alterations (i.e. neuroinflammation, autophagy dysregulation, etc.) were already established (48). To selectively knockdown neuronal NOX4, we injected AAV serotype 9 particles, which exhibits neuronal tropism for transgene delivery (61), carrying a short hairpin RNA (shRNA) targeting NOX4 (AAV-shNOX4) or a control scramble shRNA (AAV-shSCR) in the right hemispherés hippocampus (Figure 5B). AAVs encoding scramble shRNA and shRNA against NOX4 contained an enhanced green fluorescent protein (GFP) which enabled us to visualize their expression. In the AAV-shNOX4 injected mice, while GFP fluorescent signal was predominantly observed in hippocampal neurons, a total absence of signal was detected in both microglia and astrocytes, indicating that the expression of the shRNA targeting NOX4 was restricted to neuronal compartment (Supplemental Figure 7, A and B). The injection of AAV-shNOX4 decreased by 65 % mRNA levels of NOX4 to values that were similar to those observed in AAV-GFP control injected mice (Supplemental Figure 7C). Under these experimental conditions, neuronal knockdown of NOX4 was sufficient to reduce by 55% the accumulation of AT8 tau in the ipsilateral hippocampus of AAV-hTau injected mice (Figure 5, C and D). Additionally, we measured the SI and SS forms of hyperphosphorylated tau. The injection of AAV-shNOX4 7 days post AAV-hTau ICV surgery, significantly reduced AT8 SI and SS oligomers. No significant changes were observed in AT8 SI or SS monomers (Figure 5, E-G). Although, no appreciable changes were observed in either AT180 SS oligomers or monomers, neuronal NOX4 knockdown significantly decreased AT180 SI oligomers and monomers (Figure 5, H-J). These results evidence that in the AAV-hTau injected mice, despite tauopathy and its alterations were already established, neuronal-targeted NOX4 knockdown was sufficient to reduce the accumulation of pathological hyperphosphorylated tau, and mainly, its oligomeric pathological form.

**Figure 7:**
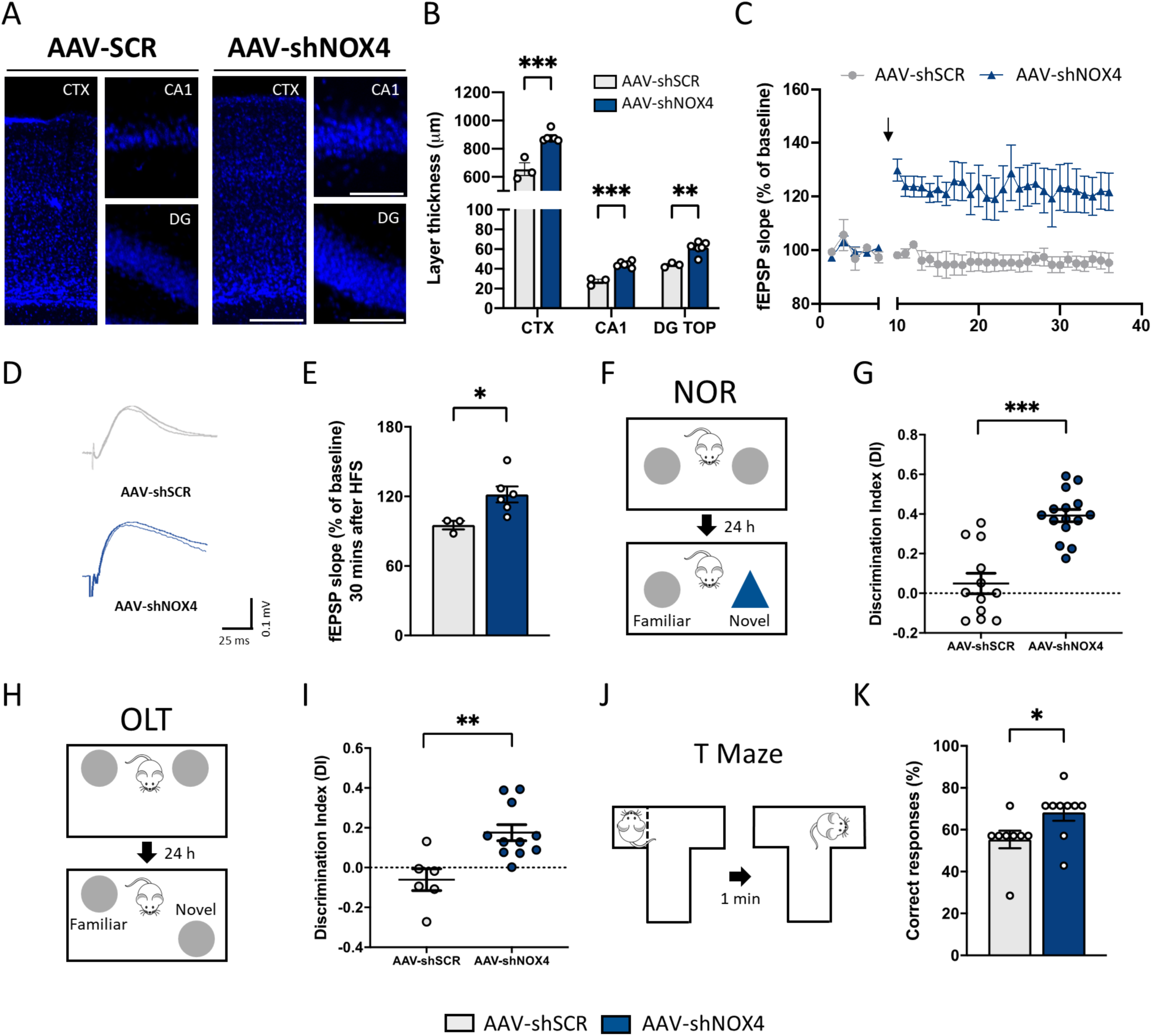
Brain atrophy, LTP impairment and cognitive decline are reduced in neuronal NOX4 knockdown mice once tauopathy is initiated. Representative images **(A)** and thickness quantification **(B)** of CTX, CA1, and DG TOP in AAV-hTau (AAV-shSCR, n=3 and AAV-shNOX4, n=6) mice. Scale bars: 100 μm (10X); 75 μm (40X). (C) *In vivo* long-term potentiation (LTP) in AAV-shSCR and AAV-shNOX4 injected mice over 30 minutes. Arrow indicates high-frequency stimulation (HFS). Five minutes of control period and 30 minutes after HFS stimulation are shown. **(D)** Representative traces before and after HFS. **(E)** fEPSP slope 30 minutes after HFS in AAV-hTau (AAV-shSCR, n=3 and AAV-shNOX4, n=6) mice. **(F)** Schematic representation of the NOR test. **(G)** Quantification of the DI in AAV-hTau (AAV-shSCR, n=12 and AAV-shNOX4, n=15) injected mice. **(H)** Schematic representation of the OLT. **(I)** Quantification of the DI in AAV-hTau (AAV-shSCR, n=6 and AAV-shNOX4, n=11) mice. **(J)** Schematic representation of the T Maze. **(K)** Quantification of the percentage of correct responses in AAV-hTau (AAV-shSCR, n=8 and AAV-shNOX4, n=9) mice. Data are presented as mean ± SEM. Significance was determined by unpaired Student’s t test or Mann-Whitney test for nonparametric data sets. *p<0.05; **p<0.01; ***p<0.001. CTX (Retrosplenial cortex); DG TOP (Dentate gyrus top layer).

### Neuronal knockdown of NOX4 modulates the ALP in tauopathy

With the aim of evaluating whether NOX4 knockdown in neurons was sufficient to prevent the macroautophagy blockade induced by hTau P301L injection (as previously observed in NOX4^-/-^ mice) we followed the experimental protocol described in Figure 5B. The macroautophagy markers p62 (Figure 6, A and B) and LC3 (Figure 6, C and D) were significantly decreased in CA1 neurons of AAV-shNOX4 injected mice compared to AAV-shSCR controls. These results, were corroborated by western blot analysis of hippocampal homogenates, where reduction in p62 (Figure 6, E-F) and LC3 II (Figure 6, E-G) was detected, indicating less accumulation of these macroautophagy markers upon neuronal NOX4 knockdown. Moreover, we observed a significant enhancement of nuclear TFEB intensity in neurons of CA1 (Figure 6, H and I) together with an increased immunoreactivity against LAMP1 (Figure 6, J and K) in AAV-shNOX4 injected mice. These results, were corroborated by western blot analysis of hippocampal homogenates; LAMP1 (Figure 6, E and L) and CTSD (Figure 6, E and M) were significantly increased in the AAV-shNOX4 injected mice, suggesting an upregulation of the lysosomal system. These results corroborate those observed in the NOX4^-/-^ mice and highlights a key role of neuronal NOX4 knockdown in preventing macroautophagy failure even when tauopathy is initiated.

### Neuronal NOX4 knockdown is sufficient to alleviate functional and cognitive impairments in tauopathy

To determine whether knockdown of neuronal NOX4 affects tau-related neurotoxicity, we measured the thickness of CTX, CA1 and DG TOP. Neuronal-targeted NOX4 knockdown significantly decreased the brain atrophy observed in AAV-shSCR injected mice (Figure 7, A and B). Moreover, while the AAV-hTau mice injected with AAV-shSCR control exhibited an LTP deficiency, the AAV-shNOX4 injected mice showed a sustained and significant LTP over 30 minutes (Figure 7, C-E). In addition, while in the NOR test (Figure 7F), AAV-hTau mice injected with AAV-shSCR control failed to recognize the novel object over the familiar one and exhibited cognitive deficit (Figure 7G), AAV-shNOX4 injected mice displayed normal cognition as shows the significant increase in the DI (Figure 7G). In the OLT test (Figure 7H), similar results were achieved to those in the NOR test (Figure 7I). In addition, in the T maze test (Figure 7J), AAV-hTau mice injected with AAV-shSCR control failed to recognize the unexplored arm during the test time, indicating spatial cognitive deficit (Figure 7K), whereas AAV-shNOX4 injected mice showed an increase in the percentage of correct responses, indicating an improvement on spatial memory (Figure 7K). Taken together, these results corroborate those obtained in the global NOX4 knockout, and demonstrate that neuronal NOX4 knockdown is sufficient to alleviate cellular loss and prevent LTP impairment and cognitive decline in AAV-hTau injected mice, even when tauopathy is established.

### The absence of NOX4 in neurons diminishes hyperphosphorylation of tau and modulates the ALP in vitro

To support the results obtained *in vivo*, primary neurons from NOX4^+/+^ and NOX4^-/-^ mice were cultured and maintained during 22 days. At day 14, neurons were subjected to PBS (as control) or AAV-hTau (to reproduce the tauopathy *in vitro*) for 8 days (Figure 8A). Under these experimental conditions, the number of AT8 positive neurons (Figure 8, B and C) and the AT8 immunoreactivity per cell (Figure 8, B and D) were significantly decreased by 54 % and 46 %, respectively, in NOX4^-/-^ neurons compared to NOX4^+/+^ neurons subjected to AAV-hTau. Several studies have demonstrated that hyperphosphorylated tau is redirected from the axonal to the somatodendritic compartment where it can impair synaptic function and cause spine loss (5, 62, 63). As Supplemental Figure 8A depicts, in neurons subjected to AAV-GFP + AAV-hTau, we found 5 different patterns of AT8 staining throughout the dendritic compartment which allowed us to classify them in a scale ranging from 0 to 4: 0) negative AT8 staining, 1) diffuse and dotted AT8 staining in the dendrites, 2) intermediate AT8 staining in the dendrites with no spine compromise, 3) intermediate to moderate AT8 staining in the dendrites with few positive spines, 4) advanced and robust AT8 staining in the dendrites with several positive spines. Remarkably, when comparing the different AT8 staining patterns with the number of dendritic spines, we found a significant negative correlation; as AT8 staining in dendrites increased, the number of dendritic spines decreased (Supplemental Figure 8B). We next studied the distribution of AT8 in NOX4^-/-^ neurons and we found a qualitative reduction in AT8 somatic missorting and a significant decrease in AT8 staining in dendrites compared to NOX4^+/+^ neurons (Supplemental Figure 8, C and D) subjected to AAV-hTau. Finally, while there is significant reduction in the number of dendritic spines in NOX4^+/+^ neurons subjected to AAV-hTau, NOX4^-/-^ neurons were protected from the dendritic spine loss induced by hTau (Supplemental Figure 8, E and F). Overall, these results indicate that NOX4 genetic deletion in neurons is sufficient to reduce the accumulation of pathologic hyperphosphorylated tau *in vitro* and correlate with those described *in vivo*. Moreover, the absence of NOX4 in neurons decreased the mislocalization of hyperphosphorylated tau to the somatodendritic compartment, preventing hTau-mediated dendritic spines loss.

**Figure 8:**
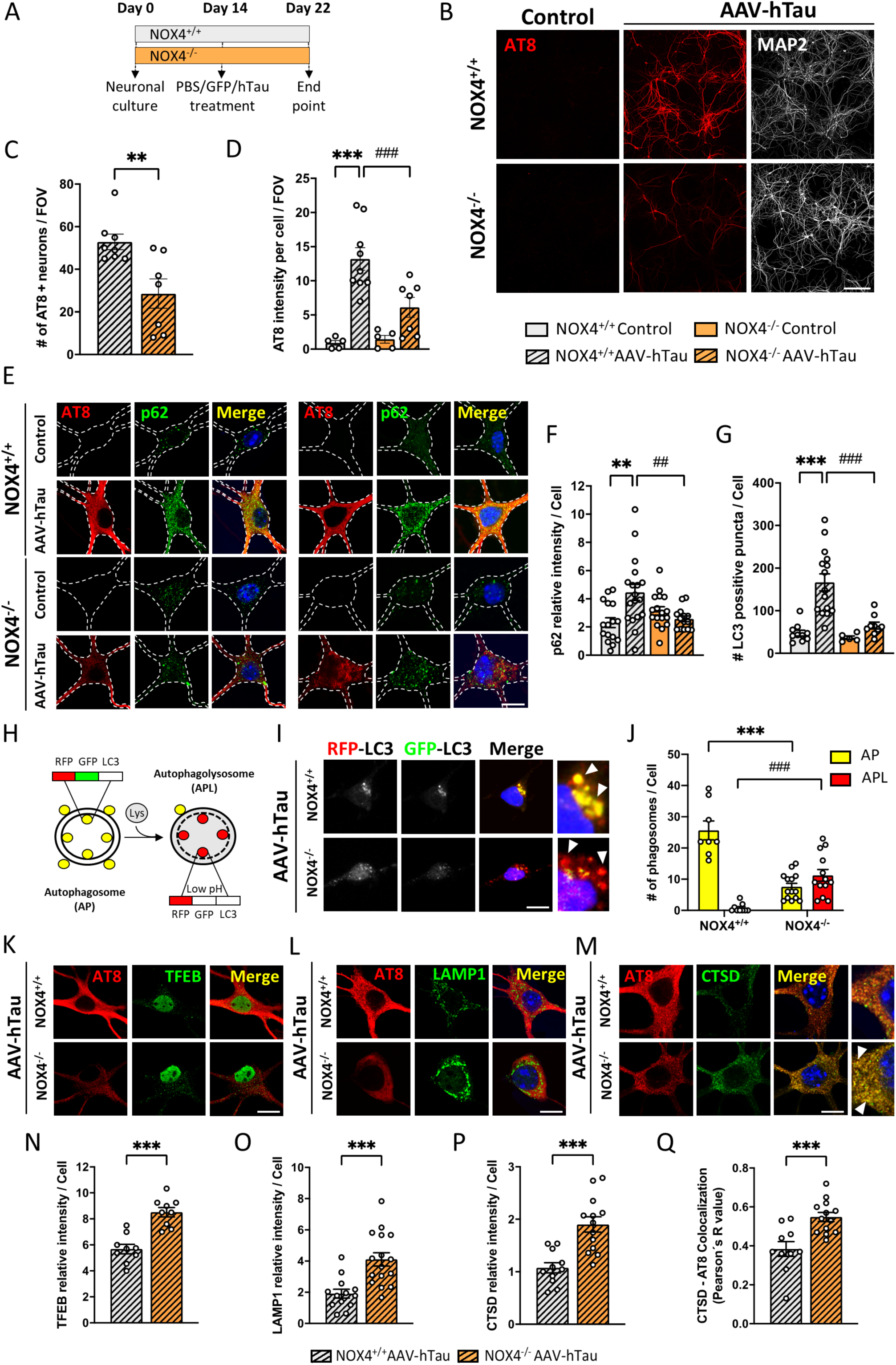
NOX4 deficiency reduces hyperphosphorylation of tau and avoid macroautophagy blockade in primary cultured neurons subjected to AAV-hTau. **(A)** Schematic representation of the protocol. Representative images **(B)** of AT8 and MAP2 relative intensity and quantification of AT8 positive neurons **(C)** and AT8 immunoreactivity per cell **(D)** in NOX4^+/+^ (Control, n=5 and AAV-hTau, n=8-9) and NOX4^-/-^ (Control, n=5 and AAV-hTau, n=7) primary cultured neurons. Scale bar: 100 μm (20X). n=Field of view analyzed per condition from 3 independent experiments. Representative images **(E)** and quantification of p62 relative intensity **(F)** and LC3 positive puncta **(G)** in NOX4^+/+^ (Control, n=10-16 and AAV-hTau, n=15-18) and NOX4^-/-^ (Control, n=5-17 and AAV-hTau, n=9-14) primary cultured neurons. Scale bar: 5 μm (63X). n=number of neurons analyzed from 3 independent experiments. **(H)** Schematic representation of the RFP-GFP-LC3 tandem reporter. Representative images **(I)** and quantification **(J)** of the number of autophagosomes (AP) and autophagolysosomes (APL) per cell in NOX4^+/+^ (AAV-hTau, n=8-9) and NOX4^-/-^ (AAV-hTau, n=14-15) primary cultured neurons transduced with the RFP-GFP-LC3 tandem reporter. Insets show images at a higher magnification. Arrows indicate autophagic vacuoles. Scale bar: 5 μm (63X). n=number of neurons analyzed from 3 independent experiments. Representative images of nuclear TFEB **(K)**, LAMP1 **(L)** and CTSD **(M)** relative intensity in NOX4 ^-/-^ and NOX4^+/+^ neurons subjected to AAV-hTau. Insets show images at higher magnification. Arrows indicate colocalization. Scale bar: 5 μm (63X). Quantification of nuclear TFEB **(N)**, LAMP1 **(O)** and CTSD **(P)** relative intensity in NOX4^+/+^ (AAV-hTau, n=9-13) and NOX4^-/-^ (AAV-hTau, n=9-16) primary cultured neurons. **(Q)** Quantification of CTSD - AT8 Pearsońs R value in NOX4^+/+^ (AAV-hTau, n=10) and NOX4^-/-^ (AAV-hTau, n=13) primary cultured neurons. n=number of neurons analyzed per each condition. Data are presented as mean ± SEM. Significance was determined by one-way ANOVA with Tukeýs post hoc test or unpaired Student’s t test. **p<0.01; ***p<0.001; ##p<0.01; ###p<0.001.

To evaluate the performance of macroautophagy *in vitro*, we performed immunofluorescence analysis of p62 and LC3 in primary cultured neurons. Both macroautophagy markers accumulated in NOX4^+/+^ AAV-hTau neurons compared to NOX4^+/+^ neurons treated with PBS, indicating a blockade in macroautophagy flux. In contrast, a significant decrease in the accumulation of p62 and LC3 was observed in NOX4^-/-^ AAV-hTau neurons (Figure 8, E-G). In neurons subjected to AAV-hTau treatment, macroautophagy was also monitored by transducing primary neurons with a tandem reporter consisting of recombinant RFP and GFP fused to LC3 protein 48 hours before the end point, a well-characterized method for assessing macroautophagy flux (64) (Figure 8H). We found that NOX4^+/+^ AAV-hTau neurons had a predominance of autophagosomes (AP) and fewer autophagolysosomes (APL). In NOX4^-/-^ neurons there was a significant decrease in the number of AP compared to NOX4^+/+^ neurons subjected to AAV-hTau and, in contrast, a remarkable increase in the number of APL (Figure 8, I and J), indicating an enhancement of the macroautophagy flux in the absence of NOX4. These results suggest that NOX4 deficiency in neurons might elicit the elimination of pathological hyperphosphorylated tau by facilitating the macroautophagy flux and corroborate the results obtained in the tauopathy *in vivo* model. Furthermore, there was a significant translocation of TFEB to the nuclei of NOX4^-/-^ neurons compared to NOX4^+/+^ neurons subjected to AAV-hTau (Figure 8, K and N). This finding, correlates with the significant augmented expression of LAMP1 (Figure 8, L and O) and CTSD (Figure 8, M and P). In addition, NOX4^-/-^ neurons presented a significant increase in the colocalization of CTSD with AT8 hyperphosphorylated tau (Figure 8, M and Q), indicating an enhanced presence of pathological tau in lysosomes. These observations support the potential implication of NOX4 regulating ALP in primary cultured neurons and explain, at least in part, the facilitation of macroautophagy flux observed *in vivo* and *in vitro* upon NOX4 deficiency.

## Discussion

By using postmortem brain samples of patients with tauopathies like AD and FTLD together with a humanized mouse model of tauopathy, this study demonstrates that NOX4 expression is upregulated in the presence of pathological hyperphosphorylated tau. Either global knockout or neuronal-targeted knockdown of the *Nox4* gene in mice was able to: 1) reduce the levels of pathological hyperphosphorylated tau, 2) modulate the ALP, 3) reduce ROS and inflammation and 4) prevent brain atrophy and synaptic dysfunction, which translate into prevention of cognitive decline *in vivo*.

A robust inverse correlation between the activity of NOX isoforms and cognitive decline has been reported in AD patients and in the APPxPS1 mouse model (37, 65). In human brain samples we observed that mRNA levels of NOX2, which is predominantly expressed in microglia under physiological conditions (66–68), were only increased in AD, as previously reported (69). However, mRNA levels of NOX4, whose expression is mostly neuronal (70, 71), were upregulated in both FTLD and AD patients, suggesting a correlation between NOX4 and tau pathology. In this line, NOX4 protein levels were overexpressed in the hippocampus and prefrontal cortex of FTLD and AD patients, key regions for learning and memory. Moreover, in the humanized *in vivo* tauopathy model, specific upregulation of *NOX4* mRNA was observed, whereas other NOX isoforms remained unaffected. Interestingly, increased expression of NOX4 was preferentially restricted to the neuronal compartment, which suggests that NOX4 may be upregulated in this cell type in response to hyperphosphorylated tau. These results, together with the increased expression of NOX2 reported in activated microglia surrounding Aβ-laden capillaries in patients with cerebral amyloid angiopathy (72, 73), suggest that NOX4 expression might be preferentially upregulated in neurons in response to pathologic tau forms, while NOX2 could be more related to the microglial response to amyloidopathy.

In this study, we show that the global absence of NOX4 and more interesting, its neuronal form, is sufficient to reduce hyperphosphorylated tau-induced toxicity through different mechanisms. First, NOX4 deletion reduced the levels of pathological hyperphosphorylated tau oligomers, which were significantly increased in SI fractions of FTLD and AD patients as well as in our *in vivo* tauopathy model. These oligomeric tau forms, which are present in the brain at early stages of AD (74, 75), have been shown to be the most toxic species in tauopathies, and are implicated in tau spreading in different cellular and *in vivo* models (76–79). Second, we showed that in NOX4 deficient primary neurons, the missorting of hyperphosphorylated tau towards the somatodendritic compartment, which causes synaptic dysfunction (5, 62, 63), was reduced, preventing the loss of dendritic spines. Third, we demonstrated that the absence of NOX4 modulates ALP, preventing the macroautophagy blockade in the *in vivo* tauopathy model. It has been reported that hyperphosphorylation of tau may alter its elimination through degradative pathways such as autophagy or the proteasome (80). In fact, the abnormal accumulation of autophagosomes in neurons of AD patients and other age-related dementias constitutes an evidence of macroautophagy failure, however the exact mechanism underlying this alteration remains unknown (1, 81). Several authors hypothesized that the build-up of autophagosomes in these NDDs may be a consequence of an enhanced autophagy induction, an impaired lysosomal degradation in later stages of the macroautophagy pathway or a coexistence of both processes (44, 82). In this line, our results indicate that neuronal overexpression of human P301L tau blocks the macroautophagy flux both *in vivo* and *in vitro*, as indicated by abnormal accumulation of autophagosomes in neurons. This blockade of the macroautophagy flux could explain, at least in part, the abnormal accumulation of hyperphosphorylated tau observed in the different tauopathy models, that does not occur in the absence of NOX4. These results agree with a recent study that demonstrates that the pathogenic P301L mutation inhibits tau degradation by interfering with different autophagy pathways, including macroautophagy, suggesting an interplay between pathological tau and autophagy that may determine its faulty degradation in the context of disease (49). Interestingly, we found that the absence of global NOX4 and neuronal-targeted NOX4 knockdown, prevented macroautophagy blockade, even when tau-related alterations were present. Furthermore, NOX4 deficiency increased the number of functional acidic autophagolysomes restoring the macroautophagy flux *in vitro*. These results suggest that an increase of neuronal NOX4 in tauopathy could play an active role in dysregulating macroautophagy flux and, thereby, contributing to disease progression.

In AD patients, increased induction of macroautophagy causes an overburden of failing lysosomes that leads to neuron toxicity, pinpointing the progressive decline of lysosomal clearance as a facilitator of the robust autophagy pathology and neuritic dystrophy implicated in AD pathogenesis (12, 19, 47, 83). Hereof, TFEB coordinately activates the expression of key genes that regulate lysosomal biogenesis and functionality and modulates genes required for autophagosome formation (84). Recent studies have provided evidence that increasing TFEB expression could be beneficial for the treatment of AD and other tauopathies (85, 86). In this study, we have observed that both NOX4 genetic deletion and neuronal-targeted NOX4 knockdown, augmented the nuclear localization of TFEB *in vivo,* together with the overexpression of its transcripts LAMP1 and CTSD. These findings can be interpreted as neuronal NOX4 downregulation can trigger TFEB nuclear translocation, increasing the availability of functional lysosomes to facilitate autophagosome degradation. In this regard, we also found increased colocalization of AT8 tau in CTSD positive lysosomes in NOX4^-/-^ neurons, suggesting enhanced delivery of hyperphosphorylated tau to lysosomes for degradation. These results, correlate with the reduction of hyperphosphorylated tau forms and may explain the reduced brain atrophy, synaptic dysfunction and cognitive impairments observed in the NOX4 knockout and neuronal NOX4 knockdown mice.

Related to the primary function of NOXs i.e. ROS production, there is growing evidence supporting that NOX-derived ROS are essential signals to activate autophagy (52, 53). In particular, NOX4 has consistently shown to induce the autophagy process (26, 52, 87). Our results evidence that NOX4 deficiency, reduces ROS production, prevents macroautophagy blockade and promotes lysosomal activity *in vivo*. Thus, we hypothesize that NOX4-derived ROS may participate in the overinduction of macroautophagy which, together with the toxic effect exerted by hTau P301L on this degradative mechanism, could determine the defective performance of this pathway, the excessive accumulation of autophagosomes and the faulty degradation of pathological forms of tau, in consistence with what has been described in several NDDs (1). In line with this assumption, it has been reported that NOX2-derived ROS promoted Parkinsonian phenotypes and protein accumulation by impairing autophagic flux and lysosomal activity (88).

Alternatively to our study, NOX4 has also been attributed an active role in the development and progression of other CNS disorders such as stroke (71), PD (89) and TBI (35, 36) which have augmented the interest in designing chemical compounds with selective NOX4 inhibitory properties (90). However, the lower specificity, selectivity, and poor pharmacokinetic profile to permeate the blood brain barrier (BBB) of available NOX4 inhibitors have impeded their study in more complex neurodegenerative preclinical models.

Overall, this study shows for the first time that NOX4 is overexpressed in different human tauopathies (e.g. FTLD and AD) and its genetic deletion and neuronal downregulation prevents tau pathology and cognitive deficits in a humanized tauopathy *in vivo* model. Furthermore, it unveils a complex role for NOX4 in modulating the ALP in neurons. In conclusion, this study validates NOX4 as a new and unexplored target for the treatment of tauopathies and highlights the potential clinical relevance of developing BBB-permeable specific NOX4 inhibitors for tau-mediated neurodegenerative disorders.

## Material and Methods

### Post mortem brain tissue

Hippocampal and prefrontal cortex frozen postmortem samples from FTLD and AD patients and non-demented controls were obtained from the Brain Tissue Bank of Fundación CIENN (Madrid, Spain). These samples were used in qPCR and western blot experiments. For immunohistochemistry experiments fixed hippocampal tissue in paraformaldehyde (PFA) from FTLD, AD and control subjects were obtained from the pathology department of VUmc and the Netherlands Brain Bank (Amsterdam, the Netherlands). The cases used in this study are summarized in Supplemental Table 1.

### Animal usage and care

For the *in vivo* experiments three-to five-month-old C57BL/6n NOX4^-/-^ and littermates wild-type NOX4^+/+^ male/female mice (25-30 g) were used. All mice were maintained in a conventional animal facility on a 12 h light/12 h dark cycle, with food and water *ad libitum*. Generation of NOX4 knockout mice was performed as previously described (71).

### AAV vectors

AAV vectors 2/6-SYN1-EGFP (AAV-GFP) and AAV2/6-SYN1-hTau (AAV-hTau) were produced and purified as previously described (48). AAV2/6-hTau particles contain the hTau mutation P301L (AAV-hTau) under the control of the neuron-specific synapsin-1 promoter, which overexpress the hTau protein specifically in neurons to achieve tauopathy *in vivo* and accelerate tau propagation (48). For *in vivo* neuronal NOX4 knockdown, AAV9-shRNA NOX4-EGFP and AAV9-shSCR-EGFP were purchased from VectorBuilder (Chicago, USA).

### Animal surgery

For ICV injections of AAV-GFP/hTau vectors, mice were anesthetized with 5% isoflurane in oxygen under spontaneous respiration. Subsequently, mice were placed in a stereotaxic instrument (David Kopf Instruments) and its temperature was monitored with a servo controlled rectal probe heating pad (Cibertec). An incision in the cranial midline was performed and skull was perforated at 0.6 mm posterior and 1.2 lateral to bregma on the right side with a micromanipulator and a micro drill. Afterwards, 1.01 μL of AAV-GFP (10^11^ VP/ml), or AAV-hTau (10^11^ VP/ml) were injected 2 mm below the dura mater (0.1 μL/min) by using an automatic Hamilton syringe (1701 N SYR). After each injection, syringes were kept in position for 6 minutes in order to avoid backflow. Control mice used for measuring oxidative stress markers in mouse hippocampal brain slices were injected ICV with 1.01 μL of PBS. For the *in vivo* NOX4 neuronal knockdown, 7 days post AAV-hTau ICV injection, AAV9-shRNA NOX4-EGFP/shSCR-EGFP were injected in the right hippocampus. Briefly, the skull was perforated at 1.94 mm posterior and 1.4 lateral to bregma on the right side by the use of a micro drill. Then, using an automatic Hamilton syringe, 1.01 μL of AAV9-sh RNA NOX4-EGFP (>10^12^ GC/ml) or AAV9-shSCR-EGFP (>10^12^ GC/ml) were injected 1.8 mm below the dura mater (0.1 μL/min). Syringes were kept in position for 2 minutes after each injection. After surgery, mice were housed for postoperative monitoring and kept until the end of the experiment at day 28.

### Behavioral tests

For assessing recognition memory, the NOR test was performed as previously described (60). Briefly, on habituation day (T0), mice were individually placed into an open-field box of 40 × 40 × 40 cm and allowed to freely explore for 10 minutes. 24 hours later, during the familiarization phase (T1), animals were placed in the previous box and allowed to explore two identical objects for 8 min. 24 hours later, on testing day (T2), in order to assess memory deficit, one object was replaced by a very different one (morphology, color and texture) and exploratory behavior was recorded for 8 minutes.

Due to the innate preference for novelty of the mice, mice without cognitive impairment will recognize the familiar object spending most of the time at the novel object. Calculation of DI was as follows: (time spent in new object − time spent in the old object) / (time spent in new object + time spent in the old object). OLT was assessed to evaluate spatial memory as described (58). On habituation day (T0), mice were placed in an empty box of 40 × 40 × 40 cm with visible spatial cues, and allowed to explore for 10 minutes. On training day (T1), 24 hours later, mice were returned to the previously explored box with two identical objects and allowed to freely explore for 8 minutes. During the testing day (T2), 24 hours later, mice were allowed to explore the box with one object moved to the opposite side. The preference for novelty was tested by determining the time spent in objects with novel and familiar locations. DI was calculated as previously mentioned evaluating novel and familiar locations. Objects and changes in object location were randomly determined and counterbalanced. Analysis was performed blinded by two independent observers using stop-watches. The T maze apparatus consists in two goal arms connected to a start arm to form a “T” shape, with a central partition in the upper middle of the “T” and two guillotine doors at the end of the right and left arms to confine the mice (91). Briefly, mice were placed in the start area with guillotine doors raised and the central partition placed. The mouse was confined in the chosen arm for 1 minute by quietly sliding the door down. After this period of time, the mouse was placed in the start area with the guillotine doors opened and the central partition removed. The test was performed seven consecutive times and a correct response was considered when the animal chose the arm that was not entered before.

### *In vivo* recordings of long-term potentiation (LTP)

Mice were anesthetized with urethane (1.6 g/Kg) and placed in a stereotaxic device. Body temperature was maintained at 37°C. Electrodes were placed stereotaxically according to the Paxinos and Franklin (2003) atlas. Field potentials were recorded through tungsten macroelectrodes (1 MΩ) placed at the CA1 region (A: -2.2; L: 1.5; V: 1-1.5 mm, from Bregma). Bipolar stainless-steel stimulating electrodes were aimed at the Schaffer collateral (SC) pathway of the dorsal hippocampus (A: -2.2; L: 2.5; V: 2 mm, from Bregma) to evoke CA1 responses. Field potentials were amplified (DAM80; World Precision Instruments, Florida, USA), bandpass filtered between 0.1 Hz and 1.0 kHz, and digitized at 3.0 kHz (CED 1401 with Spike 2 software; Cambridge Electronic Design). SC fibers were continuously stimulated with single pulses (50-200 µA, 0.3 ms, 0.5 Hz). LTP was evoked by theta-like burst stimulation (TBS) protocol, which consisted in three trains of stimuli (50 Hz, 200 ms duration), with a time-lag between trains of 200 ms (5 Hz; to mimic hippocampal theta activity). Field excitatory postsynaptic potentials (fEPSPs) were recorded during 20 min of control period and 30 min after a TBS. The initial slope of the fEPSP was assessed to quantify long-term changes of synaptic transmission. The response average during 1 minute was calculated and shown in figures. The mean average response during the 20 min period before the tetanic stimulation was considered as 100%.

### Primary neuronal culture and treatments

Primary neuronal cultures were prepared from P0 C57BL/6n NOX4^+/+^ and NOX4^-/-^ mouse embryos as previously described (92). Briefly, pups were sacrificed, and the brains were extracted and placed in Hank’s buffered salt solution. Meninges were removed and cortical and hippocampal tissue isolated, digested with papain (Sigma-Aldrich; diluted in Neurobasal (Invitrogen), DNase I (Sigma-Aldrich) (2 units/mL), EDTA (0.5 mM) and, activated with L-cysteine (Sigma-Aldrich) (1 mM) at 37°C. Afterwards, the tissue was mechanically dissociated in feeding medium (Supplemental Materials and Methods) and once centrifugated, the cell pellet was resuspended in feeding medium supplemented with 8% Fetal Bovine Serum (Sigma-Aldrich) and filtered through a 70 μm cell strainer (Corning). Neurons were plated onto 18 mm diameter coverslips previously treated with in borate buffer) and 1 hour after feeding, medium was replaced by fresh feeding medium. Neuronal cultures were maintained at 37°C in 5% CO_2_. Beginning on day 4 in vitro, feeding media was supplemented with 200 μM D, L-aminophosphonovalerate (APV) (Abcam), and feeding medium was repeated with 100 µM APV every 4 days. At day 14, neurons were treated with AAV-hTau or PBS as control and maintained up to 22 days in culture. Additionally, for visualization of dendritic spines, neurons were treated at day 14 with AAV-GFP. To monitor autophagosome and autophagolysosome formation, the Premo™ Autophagy Tandem Sensor RFP-GFP-LC3B Kit (Termofisher) was added to the neuronal cultures 48 hours before fixation. Neurons were then fixed at day 22 in 2 % paraformaldehyde for 15 minutes.

### Real-time quantitative Polymerase Chain Reaction (RT-qPCR)

Total RNA from human hippocampal and prefrontal cortex tissue and mouse hippocampal tissue was extracted with TRIzol reagent (Sigma-Aldrich) and 1 μg was reverse-transcribed using PrimeScriptTM RT Reagent Kit (perfect Real Time) (Takara). RT-qPCR was performed with qPCRBIO SyGreen Mix LoRox polymerase (Cultek) in a 7500 Fast Real-Time PCR System (Applied Biosystems by Life Technologies). Thermal cycling was carried out according to the manufacturer’s recommendations, and the relative expression levels were calculated using the comparative ΔΔCt method. The primers were obtained from Sigma-Aldrich, Madrid, Spain, (Supplemental Table 2).

### SI and SS fractions of hippocampal tissue

The protocol followed was as previously described with some modifications (93). Human hippocampal tissue and mouse ipsilateral hippocampi were homogenized in A Buffer (Supplemental Materials and Methods). Homogenates were centrifuged at 20.000 rpm for 20 minutes at 4°C. In order to obtain the SI fraction, pellets were resuspended in RAB buffer (Supplemental Materials and Methods). Then, samples were vortexed for 1 minute at room temperature, rotated at 4 °C overnight and centrifuged at 69.000 rpm for 30 minutes at 4°C. SS fractions were collected from the supernatants and SI fractions, from the pellets, which were resuspended in RAB buffer.

### Western blotting

Human hippocampal and prefrontal cortex tissue and mouse hippocampal tissue were lysed in 150 μL of ice-cold AKT lysis buffer (Supplemental Materials and Methods). 20 μg of SS, SI and protein extracts from human and mouse samples were resolved in SDS-PAGE transferred to Immobilon-P PVDF membranes (Millipore). Membranes were activated with methanol and blocked with 4 % bovine serum albumin (BSA) in Tris-buffered saline-Tween (TTBS) (Supplemental Materials and Methods) for 2 hours. Membranes were incubated with the primary antibodies (Supplemental Table 3). Then, membranes were washed three times with TTBS and then incubated with appropriate peroxidase-conjugated secondary antibody (1:10.000; Santa Cruz Biotechnology) for 45 minutes. Thereafter, membranes were washed thrice with TTBS and incubated with ECL Advance Western-blotting Detection Kit (GE Healthcare). Membranes were exposed using a ChemicDoC MP System (Bio-Rad Laboratories) and specific immunoreactive bands were quantified using Fiji software.

### Immunofluorescence

At final point, mice were deeply anesthetized with sodium pentobarbital and perfused through the ascending aorta with 0.9 % NaCl, followed by 30 mL of 4 % PFA in 0.1 M phosphate buffer (PB, pH 7.4). Brains were removed, postfixed in the same fixative at 4°C overnight, and cryoprotected for 2 days in 30 % sucrose. Forty-micrometer coronal slices were cut using a sliding microtome. For immunofluorescence assays, sections or fixed cells were abundantly washed with PB 0.1 M. Then, slices were blocked in PB 0.1M with 2 % Triton and 10 % goat or donkey serum for 1 hour, and incubated with the primary antibody (Supplemental Table 3) overnight at 4°C. Cultured cells were washed with PB 0.1 M. 0.1 % Triton (2 x 5 minutes each), blocked with 0.1 % Triton, 10 % goat serum and BSA for 1 hour, and incubated in 0.1 % Triton, 5 % goat serum and BSA with the primary antibody (Supplemental Table 3) overnight at 4°C. Tissue sections and cells were incubated with the appropriate secondary antibodies (Alexa Fluor 488, 546, 647; Invitrogen) for 1 hour and 30 minutes at 1:200 or 1:800, respectively, and then washed with PB 0.1 M (3 x 5 minutes each). In the second wash, Hoechst (33342, Invitrogen) (1 μg/mL) was added.

Human brain tissue sections were de-paraffinized in xylene and rehydrated in decreasing gradients of ethanol solutions. Afterwards, antigen retrieval was performed by transferring sections to sodium citrate buffer (pH 6.0) at 60°C for 20 minutes. Sections were then allowed to cool down for 15 minutes, and preincubated for 2 hours at room temperature in a blocking solution of tris-buffered saline 0.1 M containing 10 % goat serum and 0.3 % Triton. Sections were then incubated with primary antibodies (Supplemental Table 3), diluted in blocking solution overnight at 4°C, and washed and incubated with the appropriate secondary antibodies for 1 hour and 30 minutes prior to washing and mounting. Brain sections or fixed primary cell cultures were mounted and covered. All images were taken in a SP5 confocal microscope (TCS SPE; Leica) and processed and analyzed with Fiji software.

### Statistics

Unless otherwise stated, all data are presented as mean ± SEM. All statistical tests were performed with GraphPad (GP) Prism (version 8.3.0). Data were tested for normality to determine the use of non-parametric or parametric tests. Unless otherwise noted, all grouped comparisons were made by one-way ANOVA with Tukey’s correction and all pairwise comparisons by two-sided Student’s t-tests, depending on the experimental design. Statistical significance was set at *p< 0.05; **p< 0.01; ***p< 0.001 in accordance to GP style.

### Study approval

All experimental procedures involving animals were performed following the Guide for Care and Use of Laboratory Animals and were previously approved by the Institutional Ethics Committee of Universidad Autónoma de Madrid and the Comunidad Autónoma of Madrid, Spain, (PROEX 252/16) following the European Guidelines for the use and care of animals for research in accordance with the European Union Directive of 22 September 2010 (2010/63/UE) and with the Spanish Royal Decree of 1 February 2013 (53/2013). All efforts were made to minimize animal suffering and to reduce the number of animals used. The Ethics Committee of the Hospital Universitario La Paz, Madrid, Spain approved all protocols for human study in which experimental procedures were conducted (Ref: HULP PI-3380). Written informed consent was obtained from all subjects. The cases used in this study are summarized in Supplemental Table 1.

## Author contributions

EL contributed to design and conduct research experiments, to acquire and analyze data and to write the manuscript. PTA contributed to conduct research experiments, to acquire and analyze data and to write the manuscript. CFM contributed to conduct research experiments. AN, CP, NG and MDC contributed to perform experiments. CS and JB provided the AAV. PN and AMC provided technical advice. AR and JH provided patient samples. AIC and HHHWS provided mice colonies. MGL provided resources and contributed to design and write the manuscript. EL, PTA, CFM, MDC, HHHWS and MGL reviewed and edited the manuscript.

## Acknowledgements

This study was supported by the Spanish ministry of science, innovation and universities Ref. RTI2018-095793-B-I00 and the General Council for Research and Innovation of the Community of Madrid and European Structural Funds Ref. B2017/BMD-3827 – NRF24ADCM to MGL. EL has a fellowship from Fundación Tatiana Pérez de Guzmán el Bueno. PTA and CFM have fellowships from the Spanish ministry of science, innovation and universities. We would also like to thank Dolores Vallejo from the Confocal Unit of the Universidad Autónoma de Madrid and the Fundación Teófilo Hernando for its continuous support. All schemes were created with Biorender.com.

## Supplemental data

**Supplemental Table 1:**
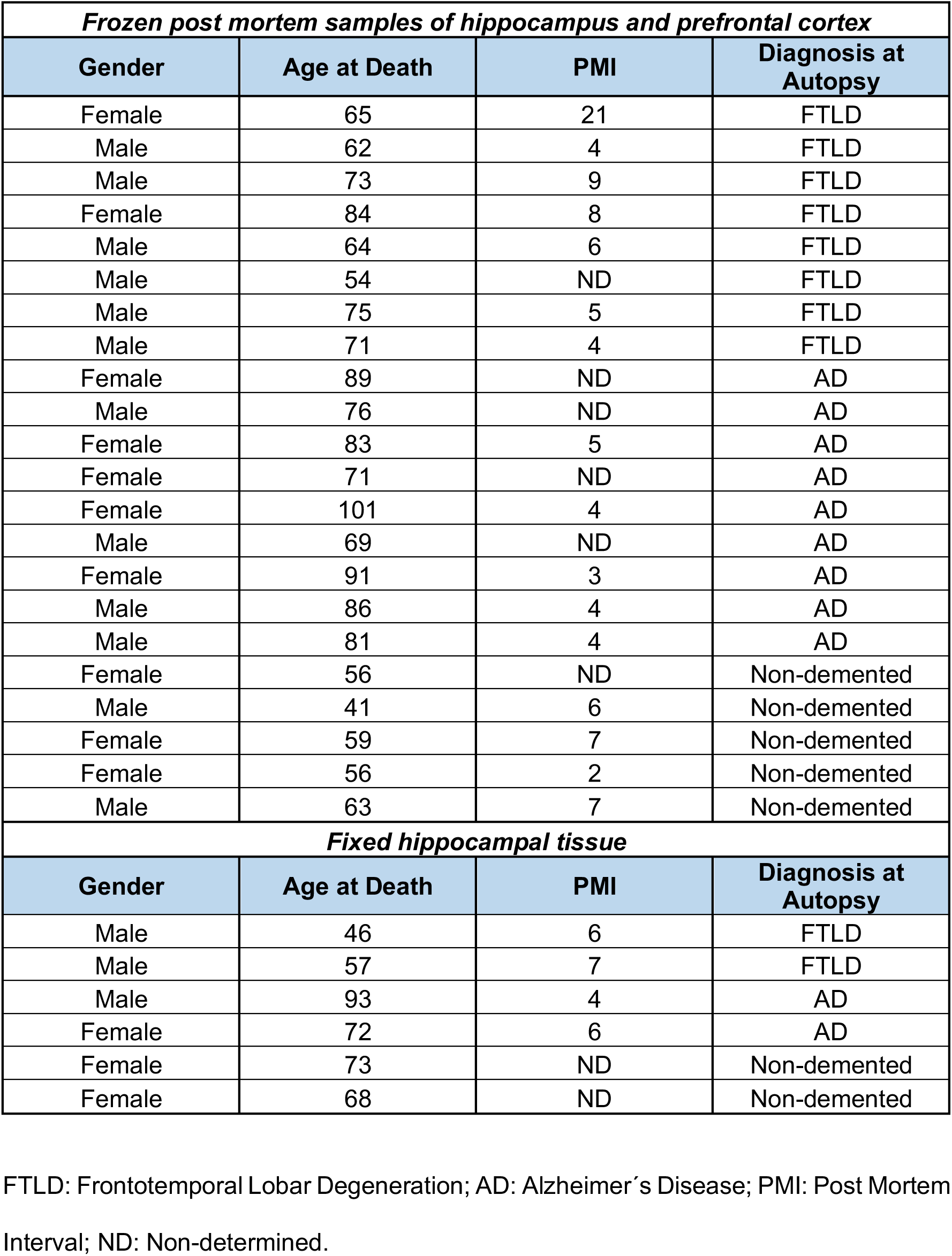
FTLD, AD and non-demented subject demographics. Related to material and method “post mortem brain tissue”.

| ***Frozen post mortem samples of hippocampus and prefrontal cortex*** |  |  |  |
| --- | --- | --- | --- |
| **Gender** | **Age at Death** | **PMI** | **Diagnosis at Autopsy** |
| Female | 65 | 21 | FTLD |
| Male | 62 | 4 | FTLD |
| Male | 73 | 9 | FTLD |
| Female | 84 | 8 | FTLD |
| Male | 64 | 6 | FTLD |
| Male | 54 | ND | FTLD |
| Male | 75 | 5 | FTLD |
| Male | 71 | 4 | FTLD |
| Female | 89 | ND | AD |
| Male | 76 | ND | AD |
| Female | 83 | 5 | AD |
| Female | 71 | ND | AD |
| Female | 101 | 4 | AD |
| Male | 69 | ND | AD |
| Female | 91 | 3 | AD |
| Male | 86 | 4 | AD |
| Male | 81 | 4 | AD |
| Female | 56 | ND | Non-demented |
| Male | 41 | 6 | Non-demented |
| Female | 59 | 7 | Non-demented |
| Female | 56 | 2 | Non-demented |
| Male | 63 | 7 | Non-demented |
| ***Fixed hippocampal tissue*** |  |  |  |
| **Gender** | **Age at Death** | **PMI** | **Diagnosis at Autopsy** |
| Male | 46 | 6 | FTLD |
| Male | 57 | 7 | FTLD |
| Male | 93 | 4 | AD |
| Female | 72 | 6 | AD |
| Female | 73 | ND | Non-demented |
| Female | 68 | ND | Non-demented |

FTLD: Frontotemporal Lobar Degeneration; AD: Alzheimer’s Disease; PMI: Post Mortem Interval; ND: Non-determined.

**Supplemental Figure 1:**
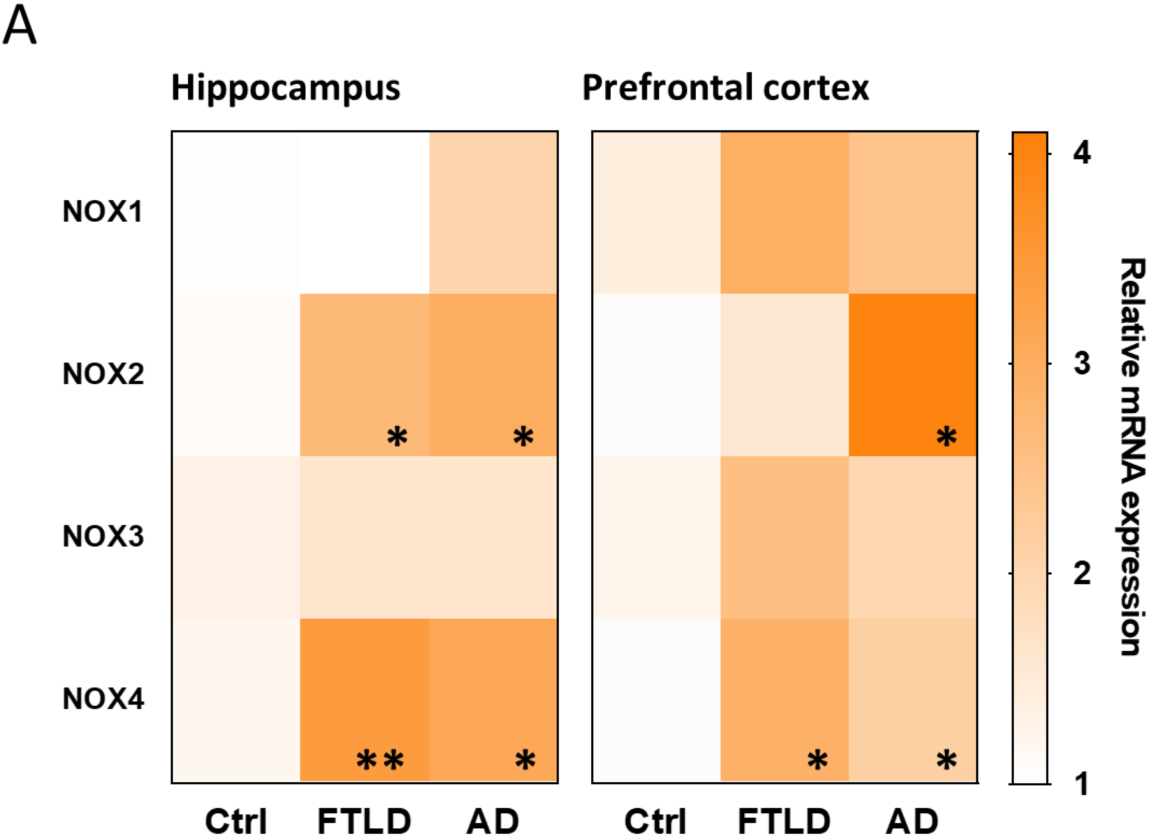
NOX isoforms transcriptional profile in FTLD, AD and non-demented subjects. **(A)** mRNA levels of NOX isoforms in human brain lysates from hippocampus and prefrontal cortex of FTLD (n=8), AD (n=9) and non-demented subjects (Ctrl) (n=5). Data are presented as mean ± SEM. Significance was determined by an unpaired Student’s t test between Ctrl and FTLD/AD. *p<0.05; **p<0.01.

**Supplemental Figure 2:**
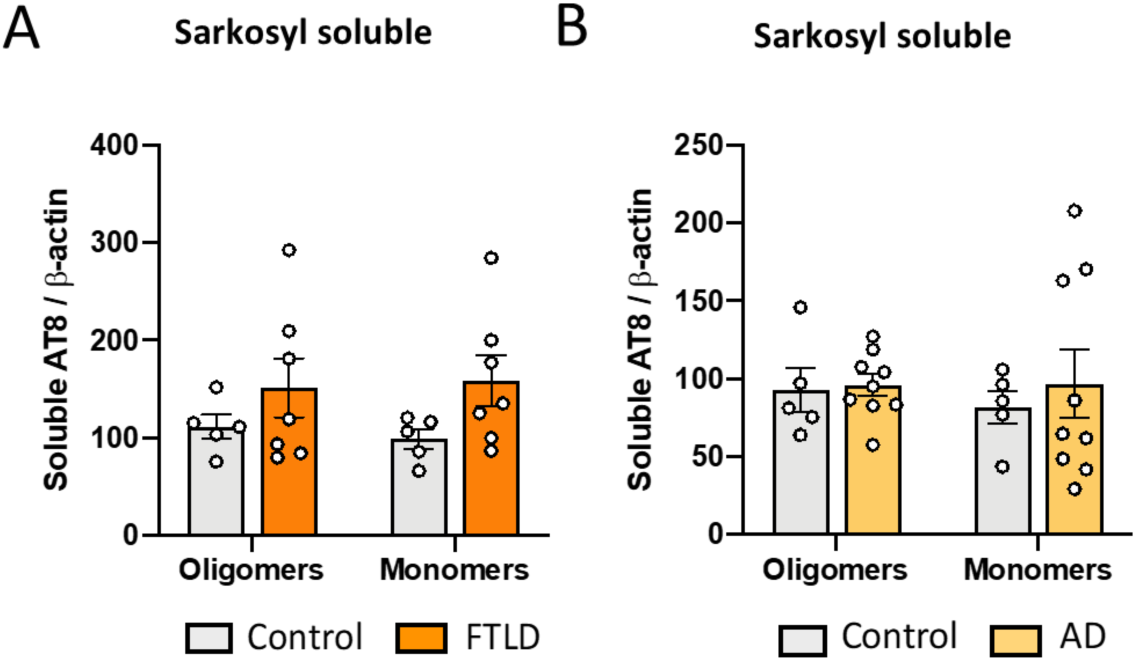
AT8 SS tau levels in FTLD, AD and non-demented subjects. **(A)** Quantification of AT8 SS tau oligomers and monomers from FTLD (n=7) and non-demented subjects (Ctrl) (n=5). **(B)** Quantification of AT8 SS tau oligomers and monomers from AD (n=9) and Ctrl subjects (n=5). Data are presented as mean ± SEM. Significance was determined by an unpaired Student’s t test. SS (sarkosyl-soluble).

**Supplemental Figure 3:**
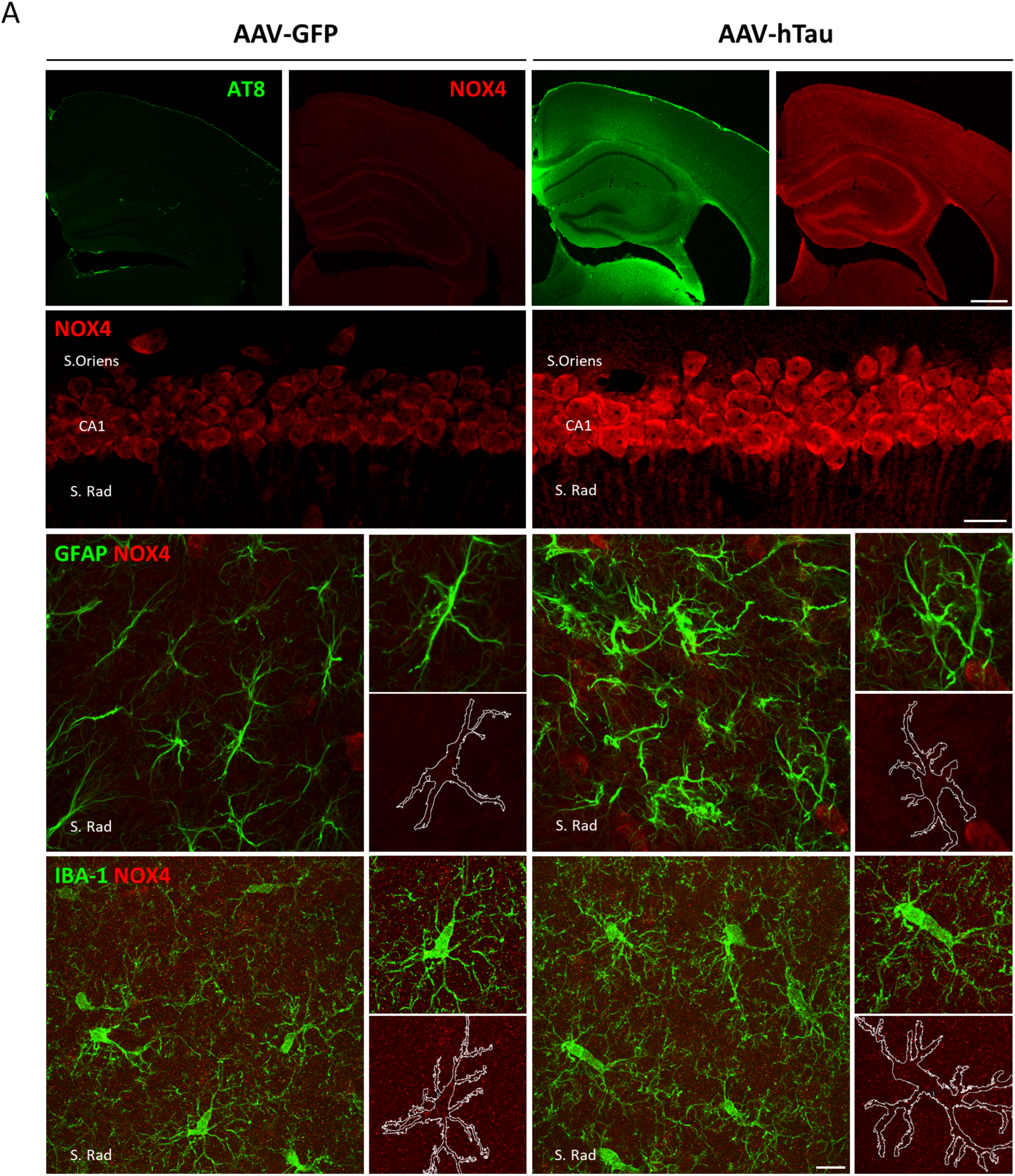
NOX4 expression in neurons and glia in the hippocampus of AAV-hTau injected mice. **(A)** Representative images of AT8 and NOX4 relative intensity in the hippocampus (upper panel), CA1 pyramidal neurons (middle panel) and glia from stratum radiatum (lower panel) in AAV-GFP and AAV-hTau injected mice. Insets show images at higher magnification. Scale bars: 500 μm (5X); 15 μm (63X). Scale bars: 10 μm (63X), respectively. S. Oriens (Stratum Oriens); S. Rad (Stratum Radiatum).

**Supplemental Figure 4:**
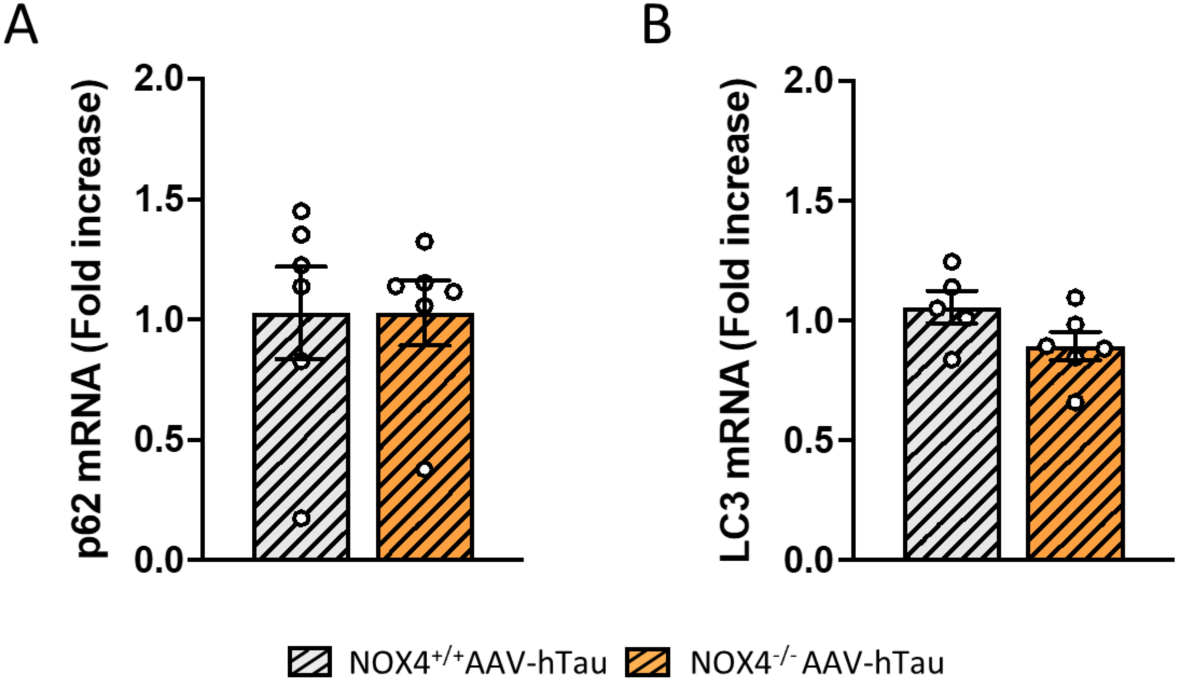
mRNA levels of p62 and LC3 in NOX4^+/+^ and NOX4^-/-^ mice injected with AAV-hTau. Quantification of p62 **(A)** and LC3 **(B)** mRNA levels in NOX4^+/+^ (n=5-6) and NOX4^-/-^ (n=6) mice injected with AAV-hTau. Data are presented as mean ± SEM. Significance was determined by one-way ANOVA with Tukeýs post hoc test.

**Supplemental Figure 5:**
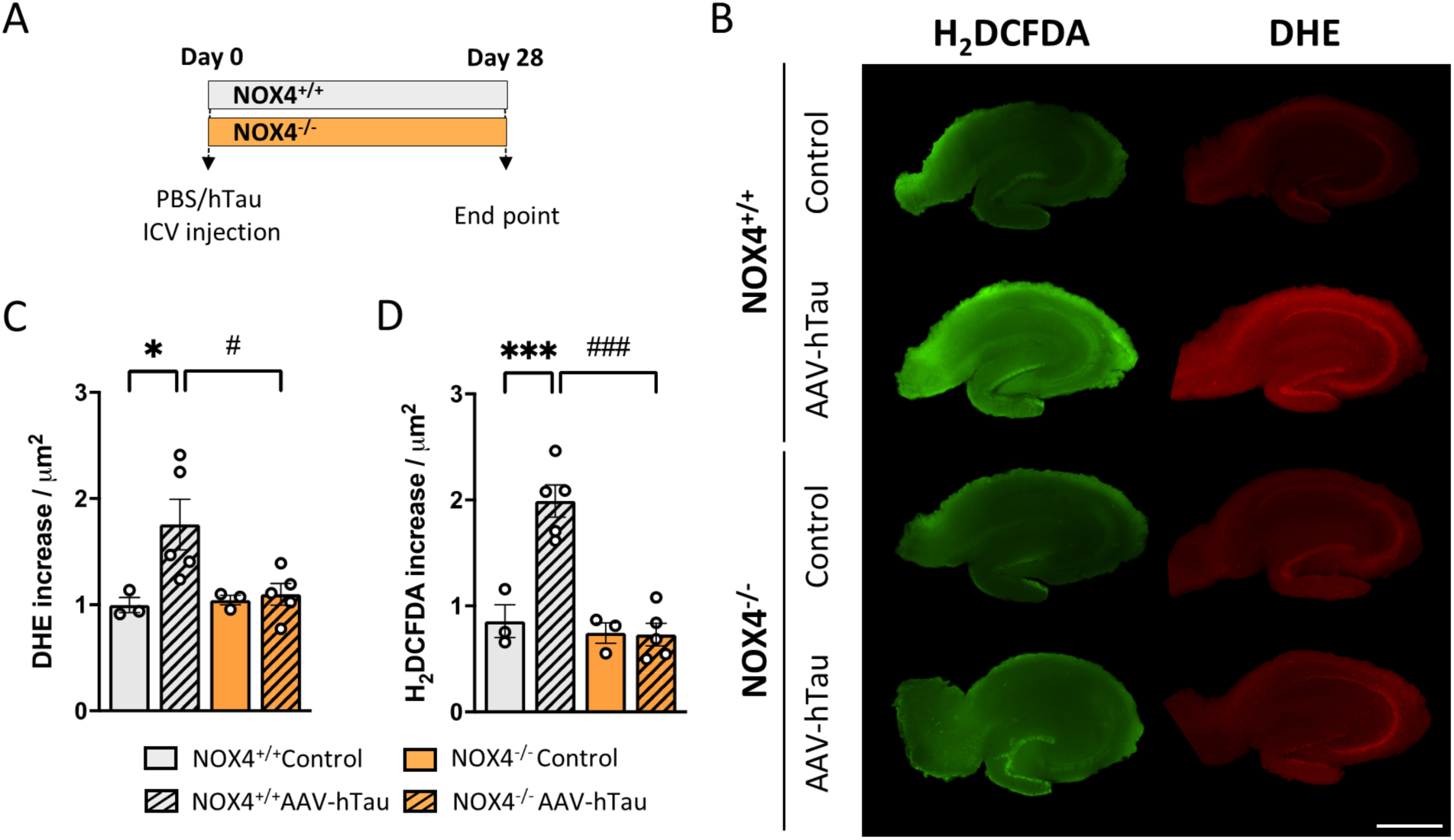
Oxidative stress markers in the hippocampus from NOX4^+/+^ and NOX4^-/-^ mice 28 days after AAV-hTau injection. **(A)** Schematic representation of the protocol. Representative images **(B)** and quantification of DHE **(C)** and H_2_DCFDA **(D)** fluorescence intensity in the hippocampus from NOX4^+/+^ (PBS, n=3 and AAV-hTau, n=5) and NOX4^-/-^ (AAV-GFP, n=3 and AAV-hTau, n=5) mice. Scale bars: 1000 μm (2X). Data are presented as mean ± SEM. Significance was determined by one-way ANOVA with Tukeýs post hoc test. *p < 0.05; ***p < 0.001; #p < 0.05; ###p < 0.001.

**Supplemental Figure 6:**
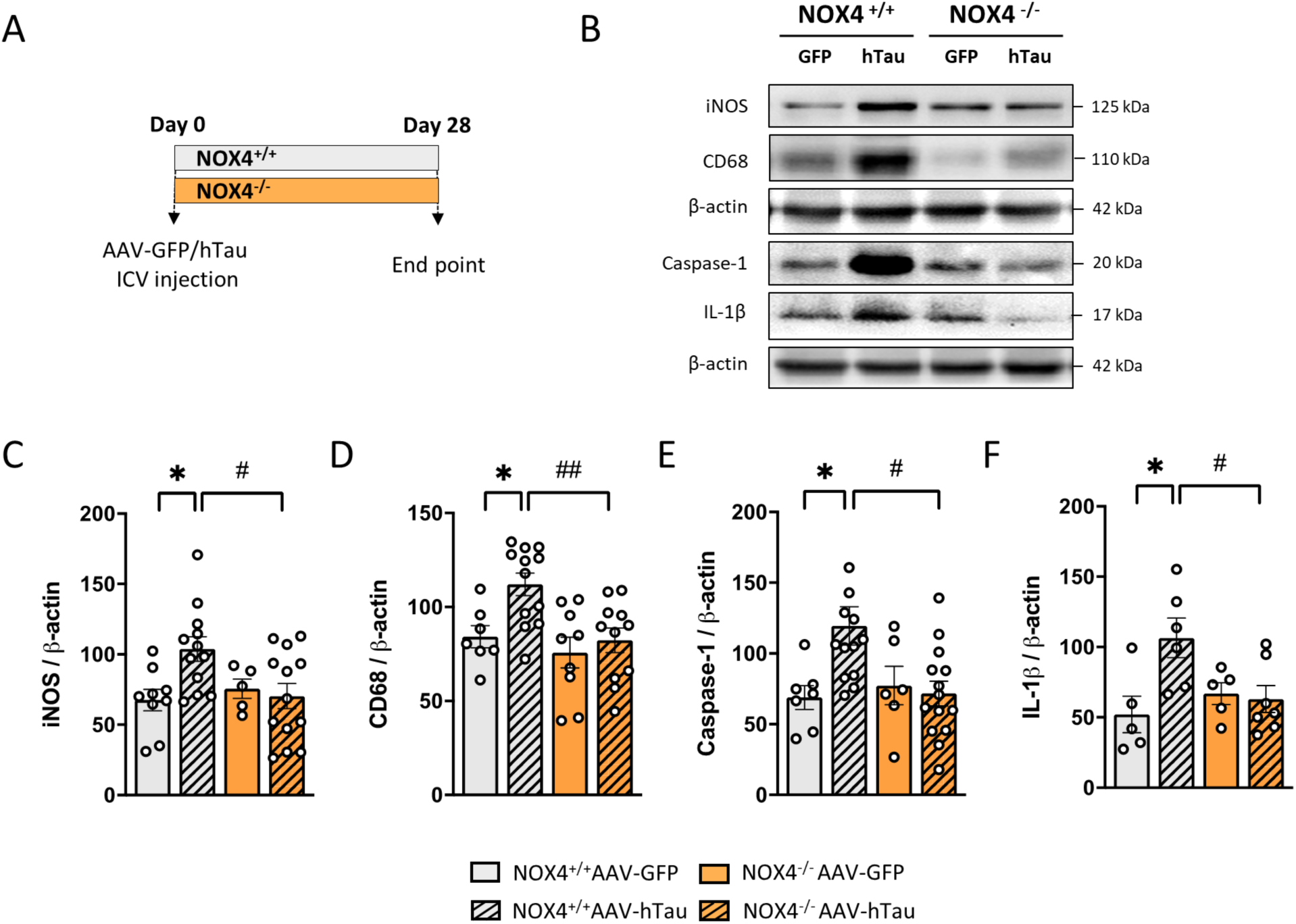
Inflammatory markers in the hippocampus of NOX4^+/+^ and NOX4^-/-^ mice 28 days after AAV-hTau injection. Representative images **(A)** and quantification **(B-E)** of protein expression levels in hippocampal lysates of iNOS, CD68, Caspase-1 and IL-1β in NOX4^+/+^ (AAV-GFP, n=5-9 and AAV-hTau, n=6-12) and NOX4^-/-^ (AAV-GFP, n=5-9 and AAV-hTau, n=7-14) mice. Data are presented as mean ± SEM. Significance was determined by one-way ANOVA with Tukeýs post hoc test. *p < 0.05; #p < 0.05; ##p < 0.01.

**Supplemental Figure 7:**
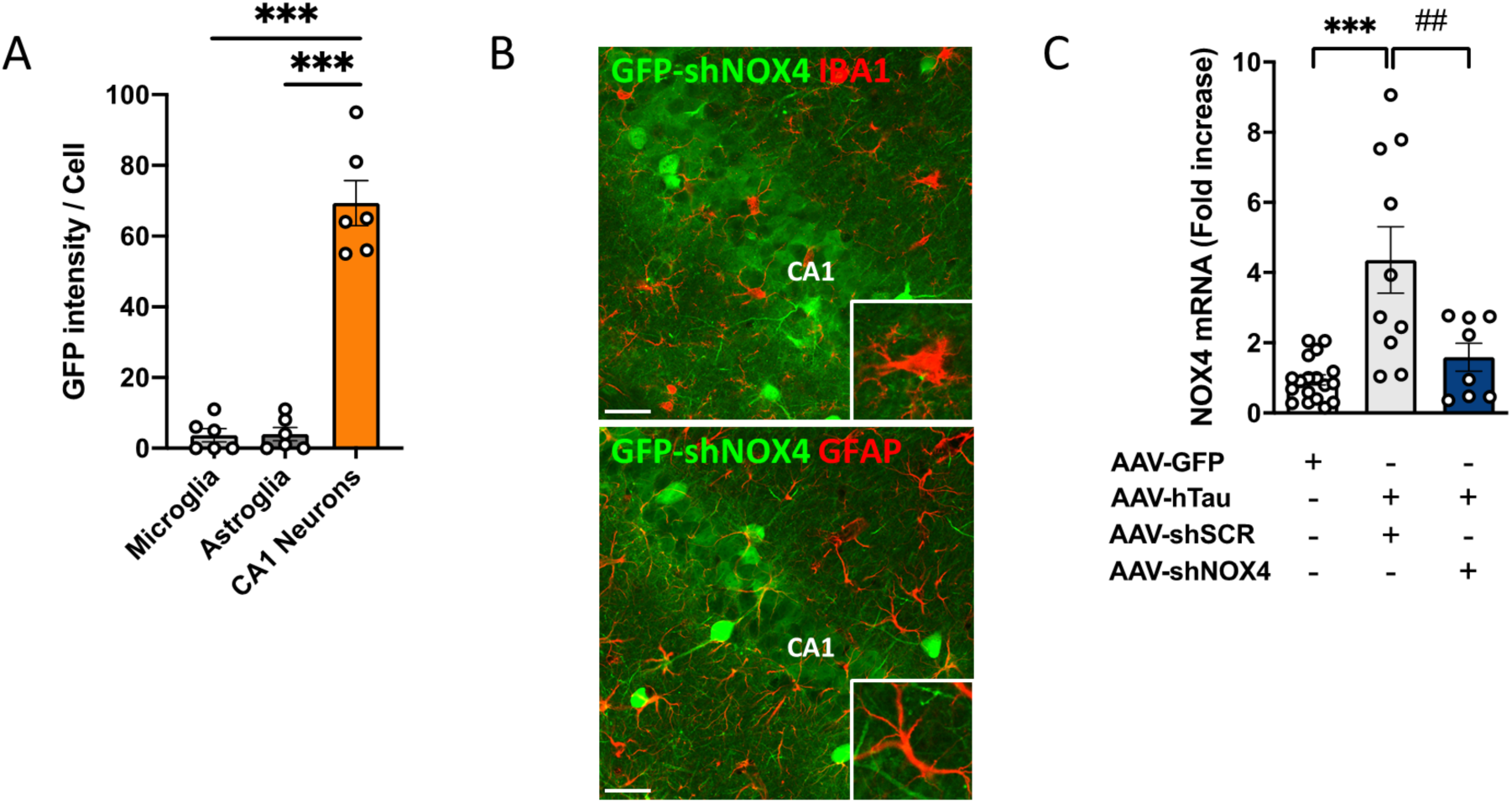
NOX4 mRNA levels in AAV-shNOX4 injected mice. Quantification **(A)** and representative images **(B)** of GFP intensity in CA1 neurons, microglia and astrocytes in AAV-shNOX4 (n=6) injected mice. Insets show images at a higher magnification. Scale bars: 25 μm (40X). **(C)** Quantification of NOX4 mRNA levels in AAV-GFP (n=18) and AAV-hTau (AAV-shSCR, n=10; AAV-shNOX4, n=8) injected mice. Data are presented as mean ± SEM. Significance was determined by one-way ANOVA with Tukeýs post hoc test. ***p < 0.001; ##p < 0.01.

**Supplemental Figure 8:**
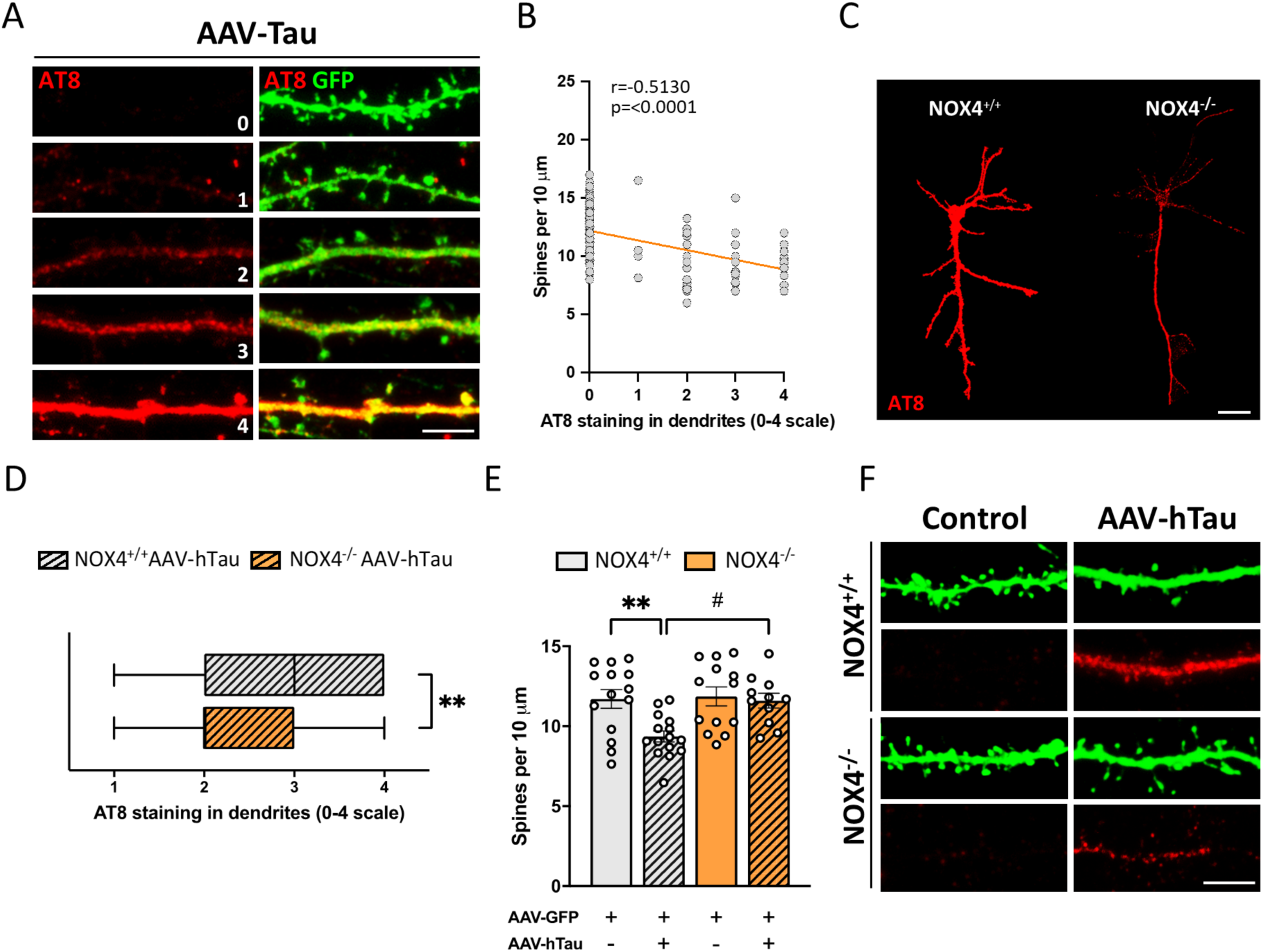
NOX4 genetic deletion diminishes the accumulation and missorting of hyperphosphorylated tau and prevents dendritic spine loss in primary cultured neurons. **(A)** Representative images of AT8 staining in dendrites in NOX4^+/+^ neurons subjected to PBS + AAV-GFP or AAV-hTau + AAV-GFP treatments. Scale bars: 5 μm (63X). **(B)** Pearsońs R value between AT8 staining in dendrites and number of dendritic spines in NOX4^+/+^ neurons subjected to PBS / AAV-hTau + AAV-GFP treatments. n=131 dendrites analyzed from 4 independent experiments. **(C)** Representative images of AT8 somatodendritic missorting in NOX4^+/+^ and NOX4^-/-^ primary cultured neurons subjected to AAV-hTau. Scale bars: 20 μm (63X). **(D)** Quantification of AT8 staining in dendrites in NOX4^+/+^ (AAV-hTau, n=70) and NOX4^-/-^ (AAV-hTau, n=58) primary cultured neurons. n=dendrites analyzed from 4 independent experiments. Quantification **(E)** and representative images **(F)** of the number of dendritic spines per 10 μm in the secondary and tertiary segments in NOX4^+/+^ (AAV-GFP, n=14 and AAV-GFP + AAV-hTau, n=15) and NOX4^-/-^ (AAV-GFP, n=13 and AAV-GFP + AAV-hTau, n=11) primary cultured neurons. Scale bars: 5 μm (63X). n=neurons analyzed from 4 independent experiments. Data represents mean ± SEM and Box and whiskers when applicable. Significance was determined by one-way ANOVA with Tukeýs post hoc test or Mann-Whitney test for nonparametric data sets. **p < 0.01; #p < 0.05.

## Supplemental materials and methods

### Evaluation of fluorescent oxidative stress markers in mouse hippocampal brain slices

28 days post injection mice were sacrificed and the hippocampi were dissected in ice-cold Krebs’s dissection buffer (Supplemental Materials and Methods). Thereafter, they were cut into 250-μm-thick slices by using a McIlwain Tissue Chopper. To allow tissue recovery after slice preparation, slices were stabilized during 45 minutes, by placing them in a preincubation solution (Supplemental Materials and Methods) pre-bubbled with 95% O_2_ / 5% CO_2_ gas mixture at 34°C. To detect the presence of ROS species as broadly as possible we used 2′,7′-dichlorofluorescein diacetate (H_2_DCFDA) and Dihydroethidium (DHE) (33). Thus, slices were incubated in control solution (Supplemental Materials and Methods) at 37 °C for 40 minutes with 10 μM H_2_DCFDA (C6827, Invitrogen), or 3.2 μM DHE (D11347, Invitrogen) in the presence of Hoechst (1 μg/mL) for 30 minutes and maintained in an incubator at 37°C in a water saturated atmosphere with 5 % CO_2_. The fluorescence was measured in a NIKON eclipse TE300 microscope (Nikon Instruments) coupled to a C9100 digital camera (Hamamatsu). Wavelengths of excitation and emission of H_2_DCFDA, DHE and Hoechst were 485, 518 or 350 and 520, 606 or 461, respectively. Hippocampal fluorescence intensity was measured using Fiji software.

### Buffers, culture media and solutions recipes

**Feeding medium:** Neurobasal + B27 plus supplement (Invitrogen) + 1x L-Glutamax (Gibco) + 1x penicillin/streptomycin (Sigma-Aldrich). **A Buffer:** 0.1 M MES Buffer pH 7.0, 1 M sucrose, 0.5 mM MgSO_4_, 1 mM EDTA, 1 mM NaF, 1 mM Na_3_VO_4_, 10 µg/ml leupeptine and phenylmethylsulfonyl fluoride (PMSF). **RAB buffer:** 0.1M MES Buffer pH 6.8, 10% sucrose, 0.5 mM MgSO_4_, 2mM EGTA, 0.5 M NaCl, 1 mM MgCl_2_, 10 mM Na_2_HPO_4_, 20mM NaF, 1 mM Na_3_VO_4_, 10 µg/ml leupeptine and PMSF, and 1% sarkosyl. AKT lysis buffer (137 mM NaCl, 20 mM NaF, 10 % glycerol, 20 mM Tris–HCl, 1 % Nonidet P-40, 1µg/mL leupeptin, 1 mM PMSF, 1 mM sodium pyrophosphate, and 1 mM Na_3_VO_4_, pH 7.5). **Tris-buffered saline-Tween** (TTBS: 10 mM Tris, 150 mM NaCl; 0.2 % Tween-20, pH 7.4). **Krebśs dissection buffer** (120 mM NaCl; 2 mM KCl; 26 mM NaHCO_3_; 1.18 mM KH_2_PO_4_; 10 mM MgSO_4_; 0.5 mM CaCl_2_; 11 mM glucose and 200 mM sucrose at pH 7.4). **Pre-incubation solution** (120 mM NaCl; 2 mM KCl; 26 mM NaHCO_3_; 1.18 mM KH_2_PO_4_; 10 mM MgSO_4_; 0.5 mM CaCl_2_ and 11 mM glucose). **Control solution** (120 mM NaCl; 2 mM KCl; 26 mM NaHCO_3_; 1.18 mM KH_2_PO_4_; 10 mM MgSO_4_; 2 mM CaCl_2_ and 11 mM glucose).

**Supplemental Table 2.**
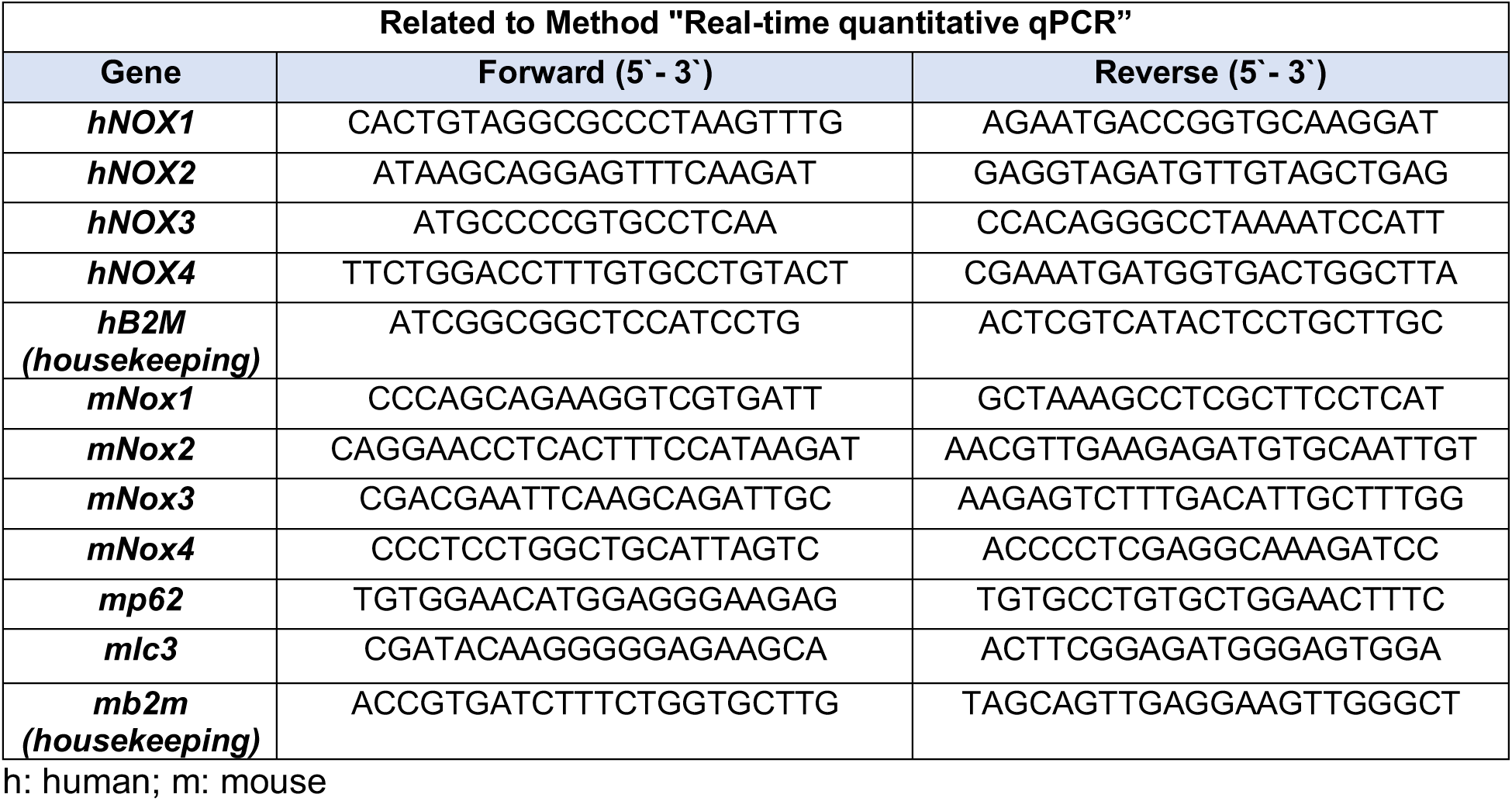
Primer sequences for qPCR.

**Supplemental Table 3.** List of antibodies used.

| Antibodies | Clone | Dilution | ID | Source |
| --- | --- | --- | --- | --- |
| **NOX4** | Rabbit monoclonal | 1:500 | Ab109225 | Abcam |
| **AT8** | Mouse monoclonal | 1:500 | MN1020 | Thermofisher |
| **AT180** | Mouse monoclonal | 1:500 | MN1040 | Thermofisher |
| **p62/SQSTM1** | Rabbit polyclonal | 1:1000 | P0067 | Sigma-Aldrich |
| **LC3A/B** | Rabbit polyclonal | 1:1000 (WB)<br>1:200 (IF) | 4108S | Cell signaling |
| **TFEB** | Rabbit polyclonal | 1:500 | 13372-1-AP | Proteintech |
| **LAMP1** | Rabbit polyclonal | 1:1000 (WB)<br>1:500 (IF) | Ab24170 | Abcam |
| **CTSD** | Rabbit monoclonal | 1:1000 (WB) | Ab75852 | Abcam |
|  |  | 1:500 (IF) |  |  |
| **GFP** | Rabbit polyclonal | 1:800 | 11-476-C100 | Exbio |
| **MAP2** | Chicken polyclonal | 1:5000 | NB300-213 | Novus Biologicals |
| **GFAP** | Mouse monoclonal | 1:1000 | MAB3402 | Sigma-Aldrich |
| **IBA-1** | Rabbit polyclonal | 1:1000 | 019-19741 | Wako |
| **iNOS** | Mouse monoclonal | 1:500 | 610432 | BD Biosciences |
| **CD68** | Rat Monoclonal | 1:500 | MCA1957GA | Bio-rad |
| **Caspase-1** | Mouse monoclonal | 1:500 | AG-20B-0042-C100 | AdipoGen |
| **IL-1β** | Hamster monoclonal | 1:500 | SC-12742 | Santa Cruz |
| **α-Tubulin** | Monoclonal | 1:10000 | T6074 | Sigma-Aldrich |
| **β-Actin** | Monoclonal | 1:50000 | A5316 | Sigma-Aldrich |

WB: Western blot; IF: Immunofluorescence

